# Epidermal clock integration and gating of brain signals guarantees skin homeostasis

**DOI:** 10.1101/2022.01.26.477844

**Authors:** Thomas Mortimer, Valentina M. Zinna, Muge Atalay, Carmelo Laudanna, Oleg Deryagin, Jacob G. Smith, Elisa García-Lara, Mireia Vaca-Dempere, Kevin B. Koronowski, Paul Petrus, Carolina M. Greco, Stephen Forrow, Paolo Sassone-Corsi, Patrick-Simon Welz, Pura Muñoz-Cánoves, Salvador Aznar Benitah

## Abstract

In mammals, an integrated network of molecular oscillators drives daily rhythms of tissue-specific homeostatic processes. This circadian clock network is required for maintaining health and is compromised by disease and lifestyle choices, such as diet and exercise. However, critical properties of this systemic network, such as which tissues communicate to coordinate their respective programs of daily physiology, and the exact homeostatic processes requiring each communication pathway, remain undefined. To dissect daily inter-tissue communication, we have constructed in mice a minimal clock network comprising only two nodes: the peripheral epidermal clock and the central brain clock. By circadian transcriptomic and functional characterization of this isolated connection, we have identified a previously unknown gatekeeping function of the peripheral tissue clock with respect to systemic inputs. That is, the epidermal clock concurrently integrates and corrects brain signals to ensure timely execution of epidermal daily physiology. Specifying the integrative arm of the clock, we identify that timely cell cycle termination in the epidermal stem cell compartment is dependent upon incorporation of clock-driven signals originating from the brain. Unexpectedly, and in contrast, the epidermal clock corrects potentially disruptive feeding-related signals to ensure that DNA replication occurs at the optimum time of day. Together, we present a novel approach for cataloguing the systemic dependencies of a given tissue, and in turn identify an essential gate-keeping function of peripheral circadian clocks that guarantees tissue homeostasis.

## Introduction

Consistent and timely execution of homeostatic programs is essential for the long-term maintenance of organismal health (*1*). In mammals, many processes that maintain cellular, tissue, and systemic homeostasis exhibit a strict 24-hour periodicity that ensures their optimal alignment to diurnal changes in the internal and external environment (*2*). To drive this daily rhythmic physiology, mammals express a 24-hour molecular oscillator (circadian clock) in almost all cell types that initiates time-of-day and tissue-specific homeostatic processes (*3*). Essential in ensuring organism-wide coherence of daily physiology is the body’s central clock, located in the hypothalamic suprachiasmatic nucleus (SCN) (*4*). The SCN clock directly receives light signals through the retina, which enables it to phase-align with external (solar) time and to subsequently drive neural, hormonal, and behavioral rhythms that synchronize peripheral clocks in other tissues. Complementing this regulatory mechanism, peripheral clocks also receive additional input directly from external cues (*5*– *7*), as well as internal signals from other peripheral clocks (*8*–*11*).

Communication between the clocks likely relies on subsets of hormones, metabolites, and neural signals, yet the exact nature of these intermediaries remains poorly defined (*12*). Furthermore, the mechanisms that allow tissue-relevant signals to be parsed from the milieu of other systemic, micro-environmental and extra-corporeal signals are unclear. Providing answers to these questions is pertinent, as incoherence introduced into this network of clocks through ageing, disease, or lifestyle changes, including diet, disrupts the daily homeostatic processes required for maintaining health (*2*, *8*, *13*–*15*). Therefore, identifying the specific function(s) of each inter-clock communication pathway is an essential first step towards understanding how this network maintains coherent daily physiology underlying health, and potentially identifying means to prevent its desynchronization in diseased states.

To date, most studies have relied on disruption of putative signaling nodes (through genetic knockout, environmental interventions, or pharmacological inhibition) to characterize the systemic inputs that control daily tissue physiology (*12*). However, these approaches cannot isolate individual clock-driven interactions between tissues, preventing comprehensive cataloguing of the daily physiology that they drive, and the mechanisms by which they do so. To circumvent this, we have reconstructed a minimal clock network *in vivo* constituting only the central clock (brain) and a single peripheral clock defined by the epidermis. This approach allowed us to dissect for the first time the specific contributions of a key signaling node (*i.e.,* the central clock) within the network in driving a tissue’s daily rhythmic physiology (*i.e*., the epidermis). In doing so, we define in unprecedented detail those rhythmic epidermal processes requiring input from the central clock and those in which the broader network of peripheral clocks (*i.e.*, non-brain clocks) are necessary. Concomitantly, and most strikingly, we show that the epidermal clock not only integrates relevant systemic signals, but also corrects those that may be detrimental to its continued homeostasis. Specifically, in proliferating epidermal stem cells, this gatekeeper function of the local clock acts to phase correct feeding-related signals that would lead to DNA replication timing that is antiphasic to physiological phase (*16*–*18*). Thus, we propose that peripheral tissue clocks act as gatekeepers to enable selective integration of the systemic signals required to drive daily tissue homeostasis.

## Results

### Tissue-specific reconstitution of BMAL1 expression isolates communication between the central (brain) clock and a peripheral (epidermal) clock

We set out to isolate (rather than to disrupt) the communication between a key signaling node and a target tissue, to understand the extent to which the node regulates the daily physiology of the tissue. We elected to focus first on signals generated by the central circadian clock in the brain, given its established role in controlling the body’s network of tissue clocks. As the peripheral tissue receiving these signals, we chose to study the intrafollicular epidermis of the skin, to exploit its extensively characterized daily physiology that becomes rewired during ageing and by different dietary conditions, such as a high fat diet or caloric restriction (*6*, *18*, *19*).

To isolate brain:epidermal clock communication, we employed our existing mouse model in which tissue-specific expression of a Cre-recombinase reconstitutes the endogenous expression of the indispensable core clock component BMAL1 - and thus clock activity - in an otherwise clock-deficient mouse (herein, *Bmal1*-StopFL mice) (*6*). Previously, we showed that keratin 14–Cre-driven reconstitution (RE) of BMAL1 expression in *Bmal1*-StopFL mice (herein, epi-RE mice) leads to clock activity and a limited daily physiology specifically in the intrafollicular epidermis—e.g. in a tissue-autonomous manner (*6*). To determine which daily physiology functions only rely on receiving brain clock signals (and do not need clock:clock communication), we generated a *Bmal1*-StopFL mouse model that expressed Cre-recombinase from the synaptotagmin 10 promoter (Syt10-Cre), which is expressed specifically in the brain (with high expression in the SCN) (*20*), to give brain-reconstituted (brain-RE) mice (Figure 1A). We have recently shown that Syt10-Cre-driven BMAL1 expression is restricted to the brain, where BMAL1 is expressed in the SCN and other brain regions such as the prefrontal cortex and hippocampus (*21*). Importantly, Syt10-Cre-driven BMAL1 expression is sufficient to drive daily cycles of activity, feeding and metabolism at the correct time of day (*21*), indicating that the SCN clock is correctly reconstituted. Finally, we combined epi-RE mice with the brain-RE mice, to produce a mouse line with reconstituted clocks in both the epidermis and the brain/SCN, giving a double-reconstituted (RE/RE) mouse model for studying brain:epidermal clock communication (Figure 1A).

**Figure 1.**
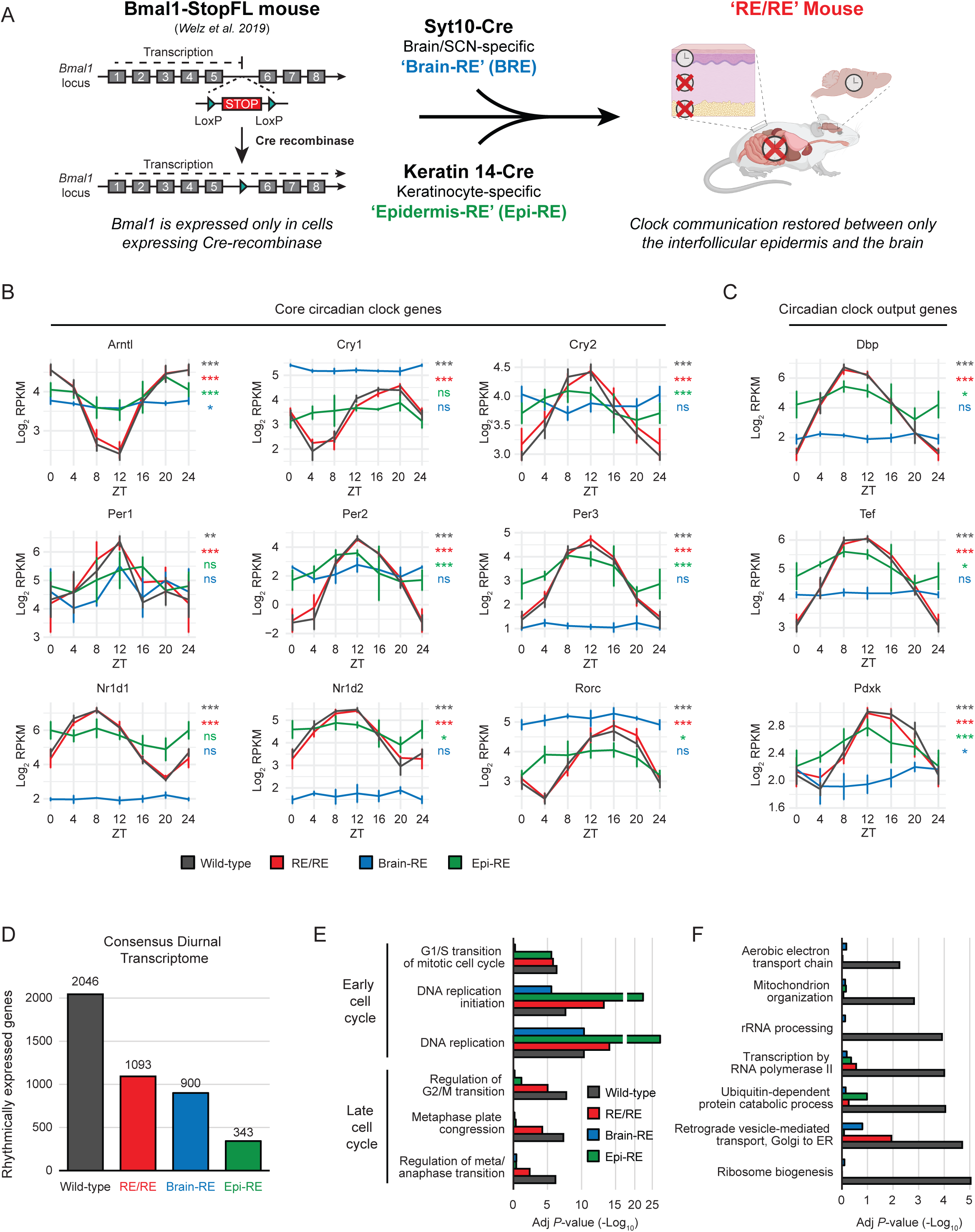
Tissue-specific reconstitution of BMAL1 expression isolates communication between the epidermal clock and brain clock: (**A**) Schematic representation of the *Bmal1*-StopFL mouse (*6*) (left) and the breeding procedure used to generate the RE/RE model (right). The image of the mouse was created using biorender.com. (**B** and **C**) Diurnal expression patterns of core clock (B) and clock-controlled genes (C) in the epidermis of all clock reconstitution conditions, as determined by RNA-seq. Values represent the log_2_-transformed mean (± standard deviation) RPKM of four biological replicates. Time point ZT0 is double-plotted for better visualization. JTK_Cycle was used to calculate the significance of each genes rhythmicity. (**D**) Quantification of the epidermal diurnal transcriptome in each clock reconstitution condition. Rhythmicity was defined as exhibiting both a JTK_CYCLE and BIO_CYCLE-calculated periodicity of 24 hours and a *P*-value ≤ 0.01. (**E**) Selected Gene Ontology terms (cell cycle–related) enriched amongst the diurnal transcriptomes of epidermis from different clock reconstitution conditions. (**F**) Selected Gene Ontology terms enriched only in the diurnal transcriptome of WT epidermis, but not other clock reconstitution conditions. Brain-RE, BMAL1 reconstituted only in brain; Epi-RE, BMAL1 reconstituted only in epidermis; RE, reconstituted; RE/RE, BMAL1 reconstituted in brain and epidermis; ns, non-significant; \**P* ≤ 0.05; \*\**P* ≤ 0.01; \*\*\**P* ≤ 0.001; RPKM, reads per kilobase per million mapped reads; SCN, suprachiasmatic nucleus; Syt10, synaptotagmin 10; ZT, Zeitgeber time.

Peripheral clocks, including those in skin, require communication with the brain clock for full entrainment to external conditions (*5*, *22*). To confirm that brain:epidermal clock communication was restored in RE/RE mice, we characterized whether the reconstituted brain clock entrained the epidermal clock. To do so, we first obtained the diurnal epidermal transcriptome from epi-RE, brain-RE, RE/RE, and WT mice (eight-week-old male mice, under a 12h:12h light/dark [LD] cycle). Strikingly, in the epidermis from RE/RE mice, genes of all core components of the epidermal clock, as well as several key clock-controlled genes, including *Dbp, Tef* and *Pdxk* (*23*–*25*), oscillated with amplitudes, phases, and periods that were indistinguishable from WT mice (Figure 1B and C). In contrast, and as previously described, oscillation patterns of core clock genes and clock-controlled genes in epidermis from epi-RE or brain-RE mice were dampened or absent, respectively, as compared to those of WT mice (Figure 1B and 1C) (*6*, *7*). Thus, we concluded that our model reconstituted known communication pathways between the brain and peripheral clocks, making it appropriate to elucidate the degree to which the brain clock controls daily epidermal physiology.

### Intact daily epidermal physiology requires communication with the brain and other peripheral tissues

Daily epidermal physiology comprises a rhythmic execution of the cell cycle, DNA repair, cell differentiation, and metabolic pathways, which together maintain tissue integrity and functionality (*2*, *17*–*19*, *26*). Underlying this daily physiology are diurnal oscillations in the expression of thousands of transcripts, which endow rhythmicity to homeostatic processes. As such, we defined high confidence diurnal transcriptomes for the epidermis of epi-RE, brain-RE, RE/RE and WT mice to determine the role of brain clock signals in driving daily epidermal physiology. To do so, we applied the rhythmicity detection algorithms JTK_Cycle and BIO_CYCLE (algorithms that use distinct methods of rhythmicity detection) (*27*, *28*) to our data sets and then generated consensus diurnal transcriptomes by intersecting the lists of rhythmic genes identified by each algorithm (Figure 1D, S1A and S1B; Supplemental table 1). In line with previous studies (*6*, *18*), 2046 genes were rhythmically expressed in WT epidermis (Figure 1D; Supplemental table 1), with a distinct night-time peak in expression (ZT18-24) (Figure S1C). Moreover, the WT epidermal diurnal transcriptome was enriched for pathways related to the mitotic cell cycle, protein homeostasis and oxidative metabolism (Figure 1E and 1F; Supplemental table 2) – all key elements of epidermal daily physiology (*6*, *18*). Unexpectedly, and in sharp contrast to the indistinguishable gene oscillations of their core clock machinery components (see: Figure 1B and 1C), the diurnal transcriptome of RE/RE epidermis constituted only 1093 genes (Figure 1D; Supplemental table 1). This RE/RE diurnal transcriptome lacked enrichment for several biological processes rhythmic in WT epidermis, such as ribosome biogenesis and mitochondrial organization (Figure 1F). However, interestingly, the RE/RE diurnal transcriptome shared with WT enrichment for pathways related to the cell cycle and vesicle transport (Figure 1E and F; Supplemental table 2). Importantly, this revealed that a significant part of the epidermal daily physiology had not been recovered despite restored epidermal:brain communication, indicating that inputs from other clock nodes are required for intact daily epidermal physiology.

Epi-RE and brain-RE epidermis also exhibited extensive diurnal transcriptomes (343 and 900 genes, respectively) (Figure 1D; Supplemental table 1). This observation agreed with the ability of the epidermal clock to drive autonomously a limited portion of the epidermal diurnal transcriptome (*6*), and the capacity of the brain clock to impose clock-independent rhythmicity upon peripheral tissues (*21*). Interestingly, the diurnal transcriptomes of epi-RE and brain-RE mice were almost exclusively enriched for genes involved in the early cell cycle (G1/S transition, DNA replication) (Figure 1E; Supplemental table 2). This contrasted with the WT and RE/RE diurnal transcriptomes where pathways related to both the late (G2/M transition, cytokinesis) and early cell cycle were present (Figure 1E; Supplemental table 2). Together, this suggested the intriguing possibility that tissue-autonomous and systemic mechanisms may drive rhythms in the early cell cycle, whilst brain:epidermal clock communication is required for physiological rhythmicity of the late cell cycle.

Direct intersection of diurnal transcriptomes does not allow statistically rigorous identification of gene sets showing equivalent or differing rhythmic expression between experimental conditions (*29*, *30*). Thus, we applied the algorithm DryR (*31*) that classifies genes into distinct sets, which share patterns in their rhythmicity parameters between multiple experimental conditions i.e. enabling us to define the genes reliant upon different clock communication pathways for their rhythmic expression. DryR identified 629 genes (BICW ≥ 0.4, Amp ≥ 0.25, Periodicity = 24 hours) rhythmically expressed with similar amplitude and phase in WT and RE/RE epidermis, but not detectably rhythmic in epi-RE or brain-RE (Figure 2A and S2A; Supplemental table 1). That is, the rhythmic expression of these genes required the communication between the brain and the epidermal clocks. Strikingly, and in line with analyses of the whole diurnal transcriptome (Figure 1E), this gene set was almost exclusively enriched for genes involved in the late cell cycle (G2/M checkpoint, mitotic spindle organization, metaphase-anaphase transition) (Figure 2B; Supplemental table 2), strongly suggesting a key role for brain:epidermal clock communication in ensuring timely cell cycle termination. This included essential regulators of the G2/M transition (CDK1, cyclin B1, and cyclin B2), and of mitotic spindle formation (Aurora kinase A, CDC20 and CENPA) (Figure 2C). Amongst brain:epidermal clock communication regulated genes, known targets of the G2/M master regulator FOXM1 (*32*) were strongly enriched (Figure 2D), and concordantly, FOXM1 was rhythmically expressed only in WT and RE/RE epidermis (Figure S2B). FOXM1 is proposed to mediate circadian regulation of the late cell cycle (*33*), and thus brain clock signals may synergize with the epidermal clock to regulate this factor. We thus concluded that correct timing of late cell cycle execution likely relies on brain:epidermal clock communication.

**Figure 2.**
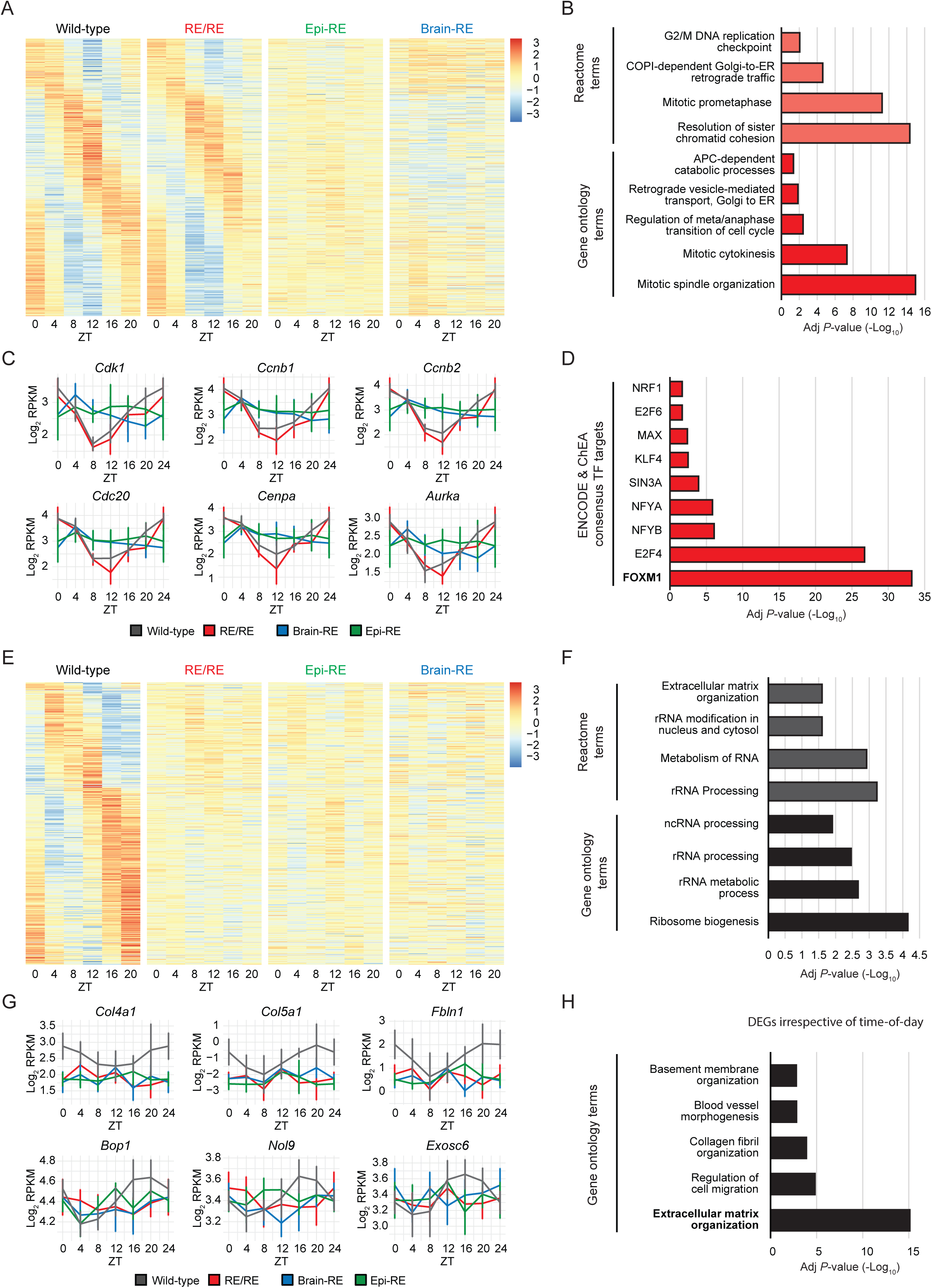
Communication with brain and other peripheral tissues ensures intact daily epidermal physiology: (**A** and **E**) Heat maps showing the diurnal expression patterns of genes equivalently rhythmic only in WT and RE/RE epidermis (629 genes) (A) or rhythmic only in WT epidermis (399 genes) (E), in all clock reconstitution conditions. Colors represent a Z-score calculated using the mean normalized counts (as calculated by DryR) from each time point. Genes are sorted by expression phase. (**B** and **F**) Selected Gene Ontology and Reactome terms enriched amongst genes with equivalent rhythmic expression only in WT and RE/RE epidermis (B) (see A) or rhythmic only in WT epidermis (F) (see E). (**C** and **G**) Diurnal expression patterns of late cell cycle regulators (C) and extracellular membrane/ribosome components (G) in the epidermis of each clock reconstitution condition, as determined by RNA-seq. Values represent the log_2_-transformed mean (± standard deviation) RPKM of four biological replicates. The time point ZT0 is double-plotted for better visualization. (**D**) Enrichment of consensus transcription factor targets (identified in ChEA and ENCODE) amongst genes equivalently rhythmic in WT and RE/RE only (see A). (**H**) Selected Gene Ontology terms enriched amongst genes differentially expressed between WT and RE/RE epidermis, irrespective of time-of-day. Differentially expressed genes were defined as those having a Padj ≤ 0.05 and fold-change ≥ 1.5. Brain-RE, BMAL1 reconstituted only in brain; ChEA, ChIP Enrichment Analysis; ENCODE, Encyclopedia of DNA Elements; Epi-RE, BMAL1 reconstituted only in epidermis; RE, reconstituted; RE/RE, BMAL1 reconstituted in brain and epidermis; RPKM, reads per kilobase per million mapped reads; ZT, Zeitgeber time.

A further 399 genes were rhythmically expressed only in WT epidermis i.e. required communication between the epidermal clock and other peripheral clocks (Figure 2E and S2C; Supplemental table 1). Contrasting with those regulated by brain:epidermal clock communication, this gene set was enriched for pathways of ribosome biogenesis and maintenance of the extracellular matrix (ECM) (Figure 2F; Supplemental table 2). This included the processing machinery of ribosomal RNAs (BOP1, EXOSC6, NOL9) and ECM proteins required to build the epidermal basement membrane (COL4A1, COL5A1, FBNL1) (Figure 2G). An intact basement membrane is required to ensure correctly executed keratinocyte differentiation (*34*), suggesting that peripheral:epidermal clock communication may guarantee faithful cell fate commitment by maintaining basement membrane homeostasis. Transcription factor (TF) binding analysis revealed significant enrichment for MYC, MAX and CREB1 targets amongst peripheral:epidermal controlled genes (Figure S2D); a set of TFs distinct to those mediating brain:epidermal communication (Figure 2D). The MYC-MAX complex negatively regulates keratinocyte expression of ECM components (*35*), and could offer a means for peripheral:epidermal clock communication to modulate ECM composition and interactions.

To establish the necessity of peripheral:epidermal clock communication to maintain epidermal homeostasis, we performed differential expression analysis comparing the transcriptomes of WT and RE/RE epidermis irrespective of time-of-day. We identified 333 differentially expressed genes (DEGs) (Padj ≤ 0.05, FC ≥ 1.5), with the majority of genes upregulated in WT relative to RE/RE epidermis (10 downregulated, 323 upregulated) (Supplemental table 1). Importantly, DEGs were most highly enriched for pathways related to assembly and maintenance of the dermal ECM (Figure 2H; Supplemental table 2), with further enrichment for pathways involved in cell motility, organ development and FGF signaling. We therefore concluded that the peripheral:epidermal communication plays an essential role in guaranteeing correct execution of epidermal daily physiology, with a particular importance in ensuring maintenance of ECM homeostasis.

### The epidermal clock selectively gates brain clock signals

In mice, epidermal DNA replication peaks during the night, ostensibly to avoid exposing the vulnerable DNA to the daytime peak of DNA-damaging oxidative conditions (*16*, *17*, *36*). Our data indicated that both the local epidermal clock and brain clock are able to independently drive rhythms in this key aspect of daily epidermal physiology (Figure 1E), thus we set out to characterize further this apparent dual regulation. In doing so, we identified a commonly rhythmic set of 119 genes (Supplemental table 1) that was highly enriched for pathways related to the early cell cycle and DNA replication (Figure 3A and 3C; Supplemental table 2). Most strikingly, although these genes were rhythmically expressed in WT, RE/RE, epi-RE and brain-RE epidermis, their expression was entirely antiphasic in brain-RE relative to the other conditions (Figure 3B). While transcript expression peaked at the beginning of night-time (ZT10-14) in WT, RE/RE, and epi-RE mice, it peaked at the onset of light hours (ZT22-2) when only the brain clock was present (brain-RE). Thus, when working alone, the brain clock appeared to drive a peak in DNA replication in proliferative epidermal stem cells that coincided with the timing of maximum oxidative metabolism in the epidermis (*16*). Critically, however, this peak was corrected by the presence of the local epidermal clock (Figure 3A and 3B). Genes exhibiting antiphasic expression included indispensable components of the DNA replication machinery, such as those encoding MCM2-4, GINS2, PCNA, and POLE (Figure 3D), and core regulators of the early cell cycle CDK2, CCNE1, E2F1 and CDC45 (Figure 3E). Importantly, comparison with existing data from Welz *et al.* (2019) showed that only 2 of the 119 genes oscillated in the epidermis of whole-body *Bmal1*-knockout mice (*Bmal1*-KO) (Figure S2E), confirming the reliance of the other genes on either local or brain clock activity for oscillatory expression. Together, this suggested a previously unidentified function of the epidermal clock whereby it is essential for gating signals originating from the brain clock, which would otherwise compromise the physiological phase of daily rhythms in DNA replication.

**Figure 3.**
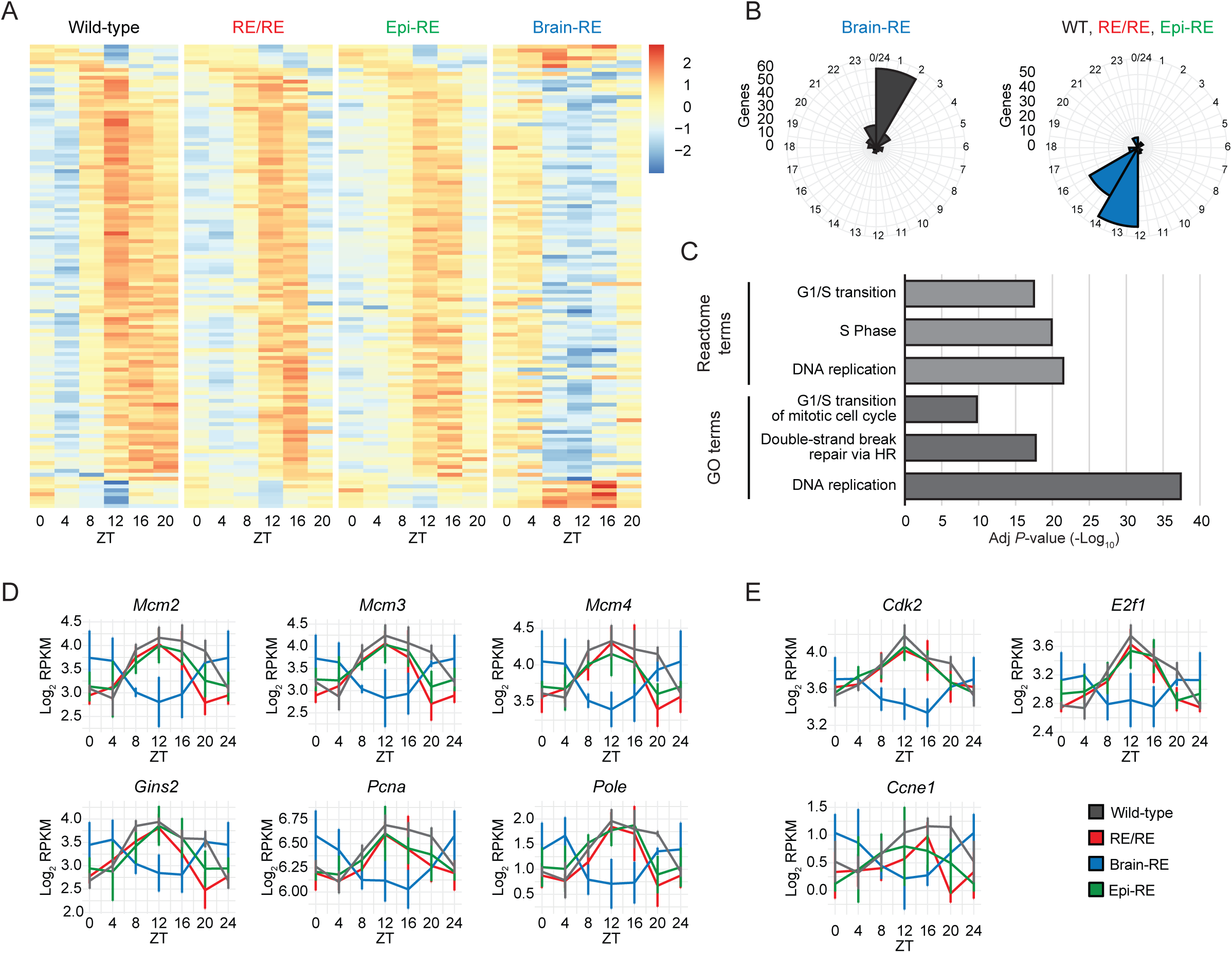
Brain clock signals drive an antiphasic pattern of epidermal DNA replication: (**A**) Heat map showing the diurnal expression patterns of genes with antiphasic rhythmicity in brain-RE epidermis relative to all other clock reconstitution conditions (119 genes), in all clock reconstitution conditions. Colors represent a Z-score calculated using the normalized read counts (as calculated by DryR) from each time point. Genes are sorted by expression phase. (**B**) Circular histograms indicating the expression phases of brain-RE antiphasic genes (see A) relative to all other clock reconstitution conditions. The phase values plotted correspond to those calculated by DryR for this model, and therefore the phase of each gene in WT, Epi-RE and RE/RE is considered equivalent. (**C**) Selected Gene Ontology and Reactome terms enriched amongst brain-RE antiphasic genes (see A). (**D** and **E**) Diurnal expression patterns of the DNA replication machinery (D) and early cell cycle regulators (E) in the epidermis of each clock reconstitution condition, as determined by RNA-seq. Values represent the log_2_-transformed mean (± standard deviation) RPKM of *n* = 4 biological replicates. The time point ZT0 is double-plotted for better visualization. Brain-RE, BMAL1 reconstituted only in brain; Epi-RE, BMAL1 reconstituted only in epidermis; RE, reconstituted; RE/RE, BMAL1 reconstituted in brain and epidermis; RPKM, reads per kilobase per million mapped reads; ZT, Zeitgeber time.

To functionally validate the apparent gating capacity of the epidermal clock, we quantified in all conditions the proportion of proliferating epidermal stem cells undertaking each phase of the cell cycle across the day by FACS analysis (8-week-old male mice, 12h:12h LD). As previously observed (*17*, *18*), WT mice showed a robust night-time peak (ZT16-20) and daytime trough (ZT4-8) of epidermal DNA replication (Figure 4A), whilst *Bmal1*-KO mice were arrhythmic (Figure 4A). In contrast, and as predicted by the transcriptional data, DNA replication rhythms in brain-RE epidermis were antiphasic (ZT4-8 peak) with respect to WT (Figure 4A). Crucially, this antiphasic pattern was corrected in RE/RE epidermis to generate WT-like rhythms of DNA replication (Figure 4A), thus confirming the ability of the local epidermal clock to gate potentially detrimental brain clock signals. The gating function of the epidermis towards brain signals with regard to DNA replication was maintained even in 35-week-old female mice (Figure S3A and S3B). To demonstrate further the capacity of the epidermal clock to correct brain signals setting the timing of DNA replication, we conditionally deleted *Bmal1* only in the epidermis to test the necessity of epidermal clock gating in the context of a fully functional clock network. Accordingly, K14-Cre-driven deletion of *Bmal1* in the epidermis led to a distinct peak in epidermal cells performing DNA replication at ZT0, representing an 8-hour phase shift with respect to WT epidermis (Figure 4C). Significantly, this further suggested that the epidermal clock is critical under physiological conditions to gate potentially disruptive brain clock signals. Overall, our results confirmed that the epidermal clock gates brain clock signals to ensure timely execution of DNA replication.

**Figure 4.**
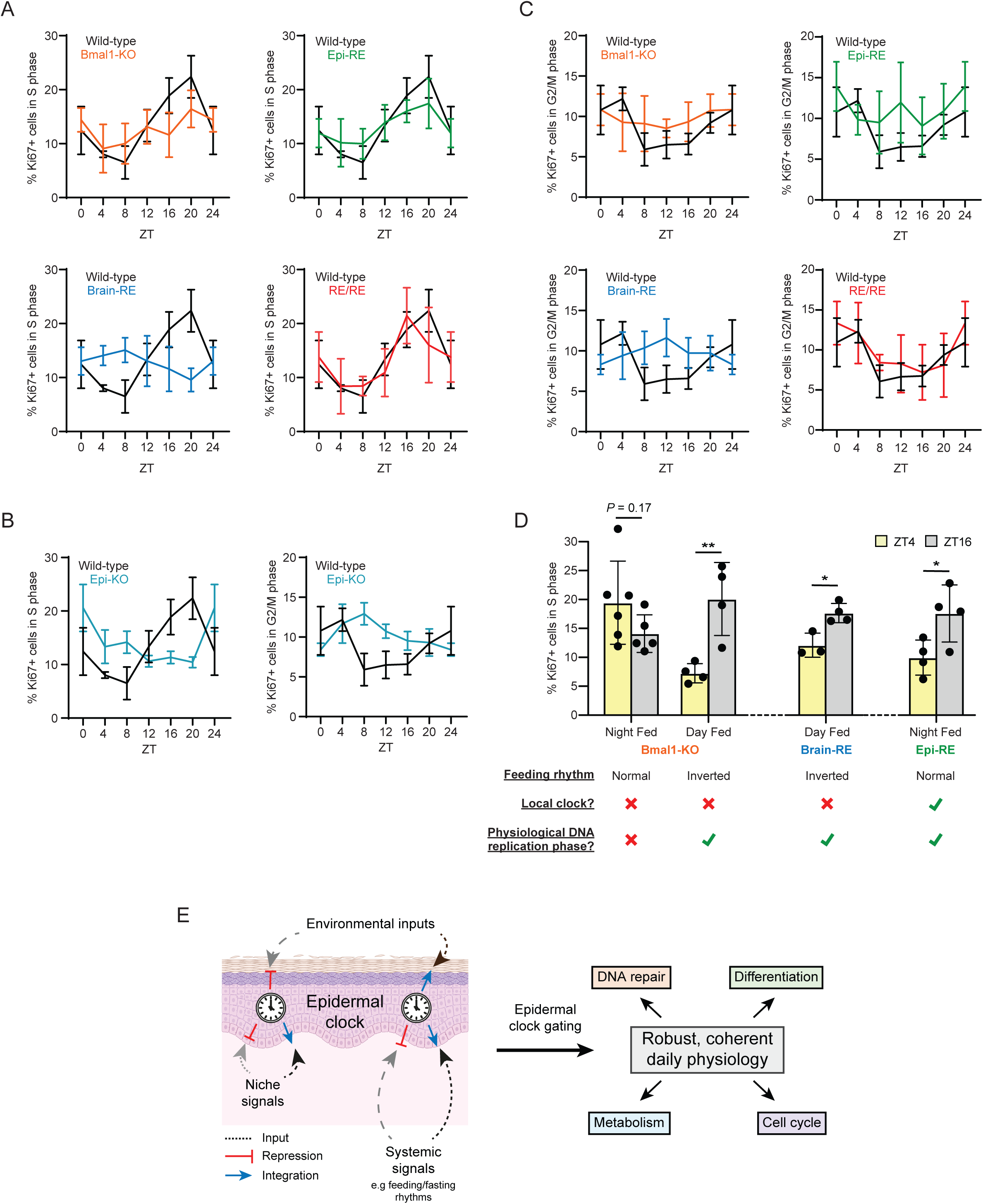
The epidermal clock corrects feeding signals to ensure physiologically phased DNA replication: (**A-C**) Percentage of actively cycling epidermal cells (Ki67-positive) in S (> 1n - < 2n DNA content) or G2/M phase (2n DNA content) across a 24-hour cycle. Values represent the mean (± standard deviation) of *n* = 4-8 biological replicates. The time point ZT0 is double-plotted for better visualization. (**D**) Percentage of actively cycling epidermal cells (Ki67-positive) in S phase (> 1n - < 2n DNA content) at ZT4 and ZT16, after two weeks exposure to different feeding regimes. Values represent the mean (± standard deviation) of *n* = 4-8 biological replicates. \**P* ≤ 0.05, \*\**P* ≤ 0.01. (**E**) Graphical representation of gating of extracellular signals by the epidermal clock. This figure was in part made using biorender.com. Ad lib, food available throughout the day; *Bmal1*-KO, full-body *Bmal1* null; Brain-RE, BMAL1 reconstituted only in brain; Day fed, food available ZT0-12; Epi-RE, BMAL1 reconstituted only in epidermis; Epi-BMAL1 KO, BMAL1 null only in epidermis; RE, reconstituted; Night fed, food available ZT12-0; ns, non-significant; RE/RE, BMAL1 reconstituted in brain and epidermis; ZT, Zeitgeber time.

Our transcriptional analyses also predicted that brain:epidermal clock communication played the contrasting function of ensuring timely execution of the late cell cycle (Figure 1E and 2B). In agreement, WT and RE/RE epidermis showed coherent rhythms in the proportion of cells present in G2/M phases (2n DNA content) (Figure 4C) and timing of mitotic division (observed as a sharp decrease in cells with a 2n DNA content from ZT4-8) (Figure 4C). This contrasted with epi-RE and brain-RE mice, which exhibited dampened and antiphasic patterns of late cell cycle execution, respectively (Figure 4C). Moreover, K14-Cre-driven deletion of *Bmal1* in the epidermis led to an antiphasic pattern of cells performing G2/M phases relative to WT mice (likely a consequence of antiphasic G1/S execution) and no apparent synchronization of cellular division (Figure 4B). We thus concluded that brain:epidermal clock communication is sufficient for ensuring timely execution of the late cell cycle.

### The epidermal clock corrects brain-driven feeding signals to gate the timing of DNA replication

Regulation of the circadian physiology from the brain to peripheral tissues may occur via direct cues (e.g., endocrine, neuronal), indirect signals resulting from body temperature rhythms and feeding-fasting cycles, or signals driving secondary molecular rhythms in other peripheral tissues. In particular, feeding-related signals instruct and synergize with the core clock machinery of peripheral tissues to guarantee timely execution of daily physiology (*10*, *37*). However, the SCN and lung clocks are refractory to changes in feeding cycles, indicating that in certain tissues mechanisms may exist to inhibit the effects of feeding-related signals (*11*, *38*). We therefore hypothesized that the epidermal clock could be gating feeding:fasting signals to prevent aberrant timing of DNA replication. If true, reinstating normal feeding cycles (i.e. night feeding) in clock-less animals should lead to antiphasic timing of DNA replication due to the lack of a corrective epidermal clock. Indeed, night feeding (2-weeks, food available ZT12-ZT0) of *Bmal1*-KO mice induced a trend towards antiphasic rhythmicity in epidermal DNA replication relative to WT (Figure 4D). In contrast, day feeding of *Bmal1*-KO mice (2-weeks, food available ZT0-ZT12) led to a night-time peak in DNA replication (Figure 4D), indicating that epidermal cell cycle initiation is generally entrainable by feeding cycles in the absence of a functional epidermal clock, but to a non-physiological phase. Importantly, day feeding of brain-RE mice corrected the antiphasic pattern of DNA replication observed under *ad libitum* conditions (Figure 4D), in line with feeding cycles serving as the brain clock signal driving antiphasic rhythms. However, most significantly, epi-RE mice, which show no daily feeding rhythms under *ad libitum* conditions (*6*), maintained a WT-like phase of DNA replication when night feeding was imposed (Figure 4D), demonstrating the capacity of the epidermal clock to gate feeding-related signals. Together, we concluded that the local epidermal clock gates brain clock-generated feeding:fasting signals to ensure timely initiation of DNA replication in proliferating epidermal stem cells (Figure 4E).

### Brain clock communication ensures robust daily epidermal physiology in the absence of environmental cues

Robustness against short-term variations in environmental conditions is essential for maintaining a consistent daily physiology. Signals from the autonomously oscillating brain clock are key for ensuring correctly timed execution of daily physiology even when environmental cues are acutely disrupted or absent (*5*, *39*). However, it is unknown whether the brain clock alone is sufficient for this stabilizing function, or if cooperation with other elements of the clock network are also required. To define the sufficiency of brain clock communication in endowing stability to daily physiology in peripheral tissues, we characterized the epidermal circadian transcriptome in WT, RE/RE, epi-RE, and brain-RE mice (all 8-week-old females) in the absence of normal light/dark (LD) cycles (i.e., after one week of constant darkness). Despite the absence of light cues, core clock genes oscillated identically in epidermis from WT and RE/RE mice in DD conditions (Figure 5A), demonstrating that communication with the brain clock alone is sufficient to ensure a stable epidermal clock entrainment. Moreover, the rhythmicity of the clock output genes *Dbp, Tef*, and *Pdxk* was indistinguishable in epidermis from WT mice and RE/RE mice, further confirming the presence of a WT-like core clock activity in RE/RE epidermis (Figure 5B). In contrast, both core clock and clock output genes lacked rhythmicity in epidermis from epi-RE mice (Figure 5A and B), in line with the observation that the autonomous epidermal clock requires light signals for its entrainment (*18*). Thus, we concluded that the brain:epidermal clock communication represents a stable and sufficient communication pathway for driving epidermal clock entrainment even in the absence of light.

**Figure 5.**
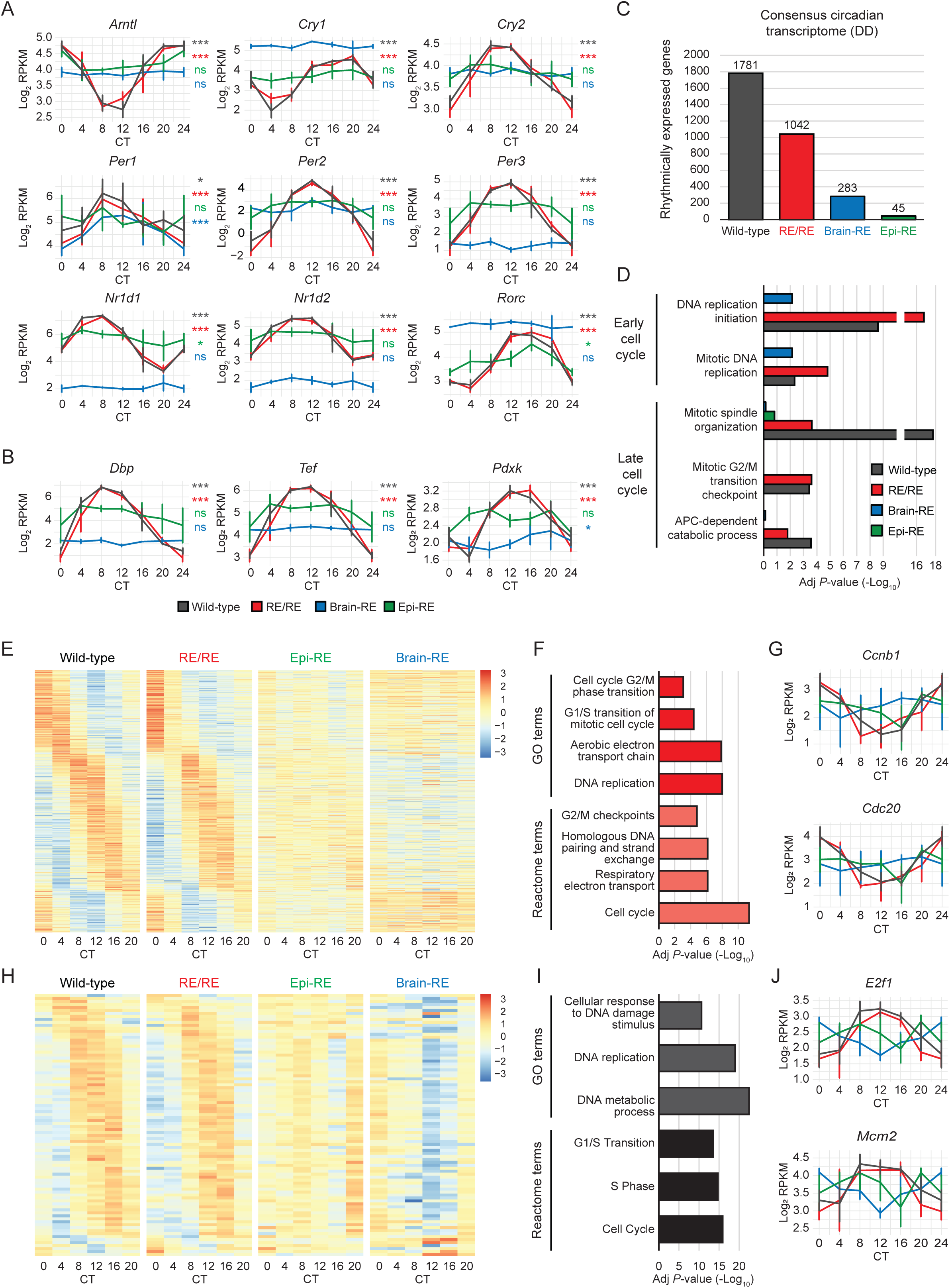
Brain clock communication ensures robust daily epidermal physiology in constant darkness: (**A** and **B**) Circadian expression patterns of core clock genes (A) and clock-controlled genes (B) in the epidermis of all clock reconstitution conditions, as determined by RNA-seq. The time point CT0 is double-plotted for better visualization. Values represent the log_2_-transformed mean (± standard deviation) RPKM of *n* = 4 biological replicates. JTK_Cycle was used to calculate the significance of each genes rhythmicity. (**C**) Quantification of the epidermal circadian transcriptome in each clock reconstitution condition. Rhythmicity was defined as a JTK_CYCLE and BIO_CYCLE-calculated periodicity of 24 hours and a *P*-value ≤ 0.01. (**D**) Selected Gene Ontology terms (cell cycle–related) enriched amongst the circadian transcriptomes of epidermis from different clock reconstitution conditions. (**E** and **H**) Heat map showing the circadian expression patterns of genes equivalently rhythmic only in WT and RE/RE epidermis (961 genes) (E) or with antiphasic rhythmicity in brain-RE epidermis relative to WT and RE/RE (89 genes), in all clock reconstitution conditions. Colors represent a Z-score calculated using the normalized read counts (as calculated by DryR) from each time point. Genes are sorted by expression phase. (**F** and **I**) Selected Gene Ontology and Reactome terms enriched amongst genes rhythmically expressed only in WT and RE/RE epidermis (F) (see E) or antiphasic in brain-RE epidermis (I) (see H). (**G** and **J**) Circadian expression patterns of the late cell cycle (G) and early cell cycle regulators (J) in the epidermis of each clock reconstitution condition, as determined by RNA-seq. Values represent the log_2_-transformed mean (± standard deviation) RPKM of *n* = 4 biological replicates. The time point CT0 is double-plotted for better visualization. Brain-RE, BMAL1 reconstituted only in brain; CT, Circadian time; Epi-RE, BMAL1 reconstituted only in epidermis; RE/RE, BMAL1 reconstituted in brain and epidermis; ns, non-significant; \**P* ≤ 0.05; \*\**P* ≤ 0.01; \*\*\**P* ≤ 0.001; RPKM, reads per kilobase per million mapped reads.

Deriving consensus circadian transcriptomes for each condition revealed that WT mice (1781 genes) and RE/RE mice (1042 genes) maintained substantial epidermal circadian transcriptomes in dark/dark (DD) conditions, whilst epi-RE mice (45 genes) and brain-RE mice (283 genes) exhibited negligible and limited rhythmicity, respectively (Figure 5C, S4A, S4B, S4C and S4D; Supplemental table 1). Importantly, the RE/RE and WT circadian transcriptomes shared an enrichment for genes related to both the early and late cell cycle (as in male mice under LD conditions) (Figure 5D; Supplemental table 2), whereas that of brain-RE mice was enriched only for the early cell cycle (Figure 5D; Supplemental table 2). As expected, no transcriptional rhythmicity of cell cycle regulators was observed in epi-RE in the absence of light (Figure 5D; Supplemental table 2) (*6*). DryR analysis identified 911 genes rhythmically expressed with similar amplitude and phase in WT and RE/RE epidermis, but arrhythmic in epi-RE or brain-RE (Figure 5E and S5A; Supplemental table 1) i.e. brain:epidermal clock communication regulated under DD conditions. Pathways enriched amongst this gene set included the late cell cycle (Figure 5F; Supplemental table 2), further suggesting that brain:epidermal clock communication maintains physiological rhythms of late cell cycle execution even in the absence of entraining light signals. Once again, this encompassed core regulators of the G2/M transition and mitotic spindle formation CCNB1-2, CDC20 and CENPE (Figure 5G and S5B).

Under DD conditions, the epidermal clock continued to gate brain clock signals to ensure timely epidermal DNA replication. DryR identified 89 genes rhythmically expressed in WT, RE/RE, and brain-RE epidermis that were highly enriched for pathways involved in the early cell cycle (Figure 5H and 5I; Supplemental table 1 and 2), but were expressed in an opposite phase in brain-RE epidermis (Figure 5H and S5C). Again, this included genes encoding core regulators of the early cell cycle, such as CDK2 and E2F1, and key components of the DNA replication machinery, such as MCM2-3, POLE, and GINS2 (Figure 5J and S5D). Together, these results demonstrated that brain clock signals regulating the epidermal cell cycle are circadian in nature, and confirmed that the local epidermal clock must both integrate and suppress distinct systemic signals to ensure correct daily rhythms of the cell cycle.

## Discussion

We have comprehensively dissected for the first time the role of a specific signaling node in defining and coordinating a distal tissue’s daily physiology. By isolating the brain:epidermal clock communication axis, we show that this connection is sufficient to drive epidermal clock entrainment as well as for specific rhythmic homeostatic processes, yet, importantly, it is insufficient to ensure full daily physiology. In particular, the brain:epidermal clock communication alone suffices for physiological rhythmicity of the cell cycle, but not for the daily rhythmicity in ribosome biogenesis or maintenance of the extracellular matrix. In other words, these additional functions require input from other peripheral clocks, and therefore communication with niche cells and other peripheral tissues is essential for attaining a complete daily physiology. In this sense, growing evidence suggests that peripheral clock communication is a conserved motif across tissue types and organ systems (*8*– *12*). For instance, recent studies suggest that extensive communication may exist between the clocks of metabolic tissues, such as the liver and skeletal muscle, as well as between distinct cell types within individual tissues (*8*, *10*, *11*). This reliance upon input from other peripheral clocks supports the proposed federated structure of the clock network (*12*), in which each clock receives and consolidates diverse inputs from the environment, the central clock of the SCN, and other peripheral clocks to determine its daily physiology. Future studies will be necessary to determine the exact identity of the peripheral tissues/niche components needed for intact epidermal daily physiology.

Directly contrasting with the systemic dependencies of epidermal daily physiology, we have also uncovered a previously unrecognized gating function of peripheral tissues towards daily brain signals. In particular, we show that the epidermal cell cycle represents a paradigm for cooperation and conflict between the brain and peripheral clocks. If unchecked by the local epidermal clock, feeding-related signals, driven by the brain clock, would induce a morning peak in DNA replication — a profile antiphasic to that of WT epidermis. However, the presence of the local clock suppresses this feeding-related signal and corrects the phase of DNA replication. This interaction suggests that the epidermal clock selectively suppresses systemic signals that would otherwise compromise the coherence of homeostatic processes. Strikingly, and in direct contrast with the early cell cycle, we also show that brain:epidermal clock communication is sufficient to ensure physiological rhythmicity in execution of late cell cycle stages (G2/M transition and cytokinesis). These distinct interactions between brain clock signals and the early and late stages of the epidermal cell cycle suggest that each phase requires coherence to distinct external events, for which the brain clock signals suffice only in the case of the late cell cycle.

The separation of peak oxidative phosphorylation and peak DNA replication in the epidermis has led to the proposal that the two processes are segregated by the circadian clock to avoid oxidative damage to replicating DNA (*16*). Nonetheless, definitive experiments functionally linking these cell states are lacking, and it remains possible that nocturnal DNA replication may favor epidermal homeostasis for other, as yet unknown, reasons. Notably, cell proliferation is highly dependent upon aerobic glycolysis to provide the building blocks necessary for faithful cell cycle execution (*40*). Thus, a night-time peak in DNA replication may plausibly anticipate an enhanced nocturnal availability of glucose, due to feeding activity, which in turn could be used for aerobic glycolysis. A misalignment of DNA replication, as in the case of brain-RE mice, would therefore be expected to slow, or even prevent, cell cycle completion, potentially compromising maintenance of the epidermal barrier. Further experiments will be required to determine the true motive behind the nocturnal timing of epidermal DNA replication.

Feeding rhythms, with their established function in entraining most peripheral clocks, are a key central clock input to peripheral tissue clocks. Emphasizing this importance, approximately 20% of liver daily physiology relies on liver clock integration of feeding signals, and feeding rhythms are sufficient for nearly 60% of the circulatory metabolite rhythms essential for inter-tissue coordination (*10*, *21*). Here, we reveal a potentially detrimental side to feeding cycles, showing their capacity to induce untimely activation of biological processes, such as the epidermal cell cycle, when not buffered/gated by peripheral clocks. Although indispensable, daily feeding rhythms may represent a recurrent stress for many tissues, which if left unchecked could lead to loss of coherence in key homeostatic processes and progressive deterioration of tissue integrity. Thus, tissues may be obliged to employ molecular buffers, such as the circadian clock, to neutralize the effects of this daily change in their external environment. The exact identity of the feeding-related signals inducing inappropriate cell cycle entry remains unclear; however, the potent cell cycle regulatory capacity of feeding-derived metabolites and insulin signaling may represent promising candidates (*41*, *42*).

Bodily tissues exist within the context of an ever-changing milieu of extracellular signals, of which only a select few are integrated by cells to maintain tissue homeostasis. Contributing to this dynamic extracellular environment, the body’s central clock produces an array of signals that exert essential tasks, such as entraining the rest of the body to changes in light; however, the myriad signals emanating from the brain clock are not universally relevant to the many tissues upon which they impinge. As such, peripheral tissues must selectively interpret and repress systemic signals originating from the brain clock (and potentially those from other peripheral clocks), depending on whether the information they transmit is relevant to the execution of that tissue’s daily physiology (e.g., epidermal DNA replication). To parse the extracellular environment, we propose that tissues employ their circadian clock as a gatekeeper to interpret, buffer and/or correct this complex array of signals (Figure 4E). Clock-mediated gate keeping could be envisaged to occur through daily regulation of cell surface receptor expression, direct competition with aberrantly activated the TFs, amongst other possibilities. Supporting such a gating function, peripheral clocks respond differentially to entraining signals, such as feeding and light, implying tissue-specific mechanisms that integrate or suppress particular brain clock signals (*11*, *38*). Moreover, the tissue clocks of liver and muscle are able to inhibit the response of specific peripheral clocks to feeding-related signals (*10*, *11*), highlighting a potential role for inter-peripheral clock communication in regulating how a tissue responds to brain clock signals. Here, we unify these findings, implicating the local clock as both a transducer and repressor of systemic signals.

The rewiring and decay of inter-tissue communication is a recurrent feature of many disease states (*8*, *43*, *44*). However, methodologies to detail exhaustively the role of systemic inputs in ensuring tissue homeostasis, and how they change during different disease states, have remained out of reach. Here, for the first time, we present a full dissection of the elements of a peripheral tissue’s daily physiology regulated by a specific distal signaling node. In doing so, we reveal a novel gatekeeper role for the epidermal clock, through which brain clock signals are policed to ensure coherence of epidermal daily physiology. We believe that these findings represent a critical shift in how circadian clock function can be understood in the context of both health and disease.

## Acknowledgements

We would like to thank the genomics unit at the CNAG for assistance with bulk RNA sequencing and Prof. Gregor Eichele for generously providing us with the Syt10-Cre mice. We would also like to thank Veronica Raker for proofreading the manuscript.

## Author contributions

TM, VMZ, PSW and SAB designed the study with critical input from all other authors. TM and VMZ performed the majority of experiments. MA and EG provided experimental assistance with the analysis of the cell cycle by FACS. TM, VMZ, CL and OD performed the data analysis of the diurnal and circadian transcriptomes. PSW and SF generated the Bmal-StopFL mice used in this study. TM and SAB wrote the manuscript, integrating feedback from all other authors.

## Data and materials availability

The datasets associated with this study have been uploaded to the Gene Expression Omnibus under the code GSE190035. All unique/stable reagents generated in this study are available from the lead contact with a completed material transfer agreement. Further information and requests for resources and reagents should be directed to and will be fulfilled by the lead contact, Salvador Aznar Benitah.

## STAR Methods

### Housing of mice and study design

All mice were bred and maintained in the animal facilities of the Barcelona Science Park (PCB), following the applicable Spanish and European Union regulations. The experimental protocols applied in this paper were approved by the Catalan Government, in line with the relevant legislation and the guidelines of the Institutional Animal Care and Use Committee (IACUC) at the PCB.

The ARRIVE guidelines for reporting and performing animal experiments (*45*) were taken into consideration during the design of all experiments. Mice were maintained in rooms at 22°C degrees with *ad libitum* access to standard chow diet, except for feeding-related experiments, where feed was available from ZT12-0 (night feeding) and ZT0-12 (day feeding). To profile daily transcriptional rhythms, male mice were maintained under standard 12h:12h light/dark conditions (LD), and four eight-week-old mice were collected per time point. To profile circadian transcription, seven-week-old female mice were maintained in constant darkness (DD) for between 156 and 176 hours and then sacrificed (at eight weeks of age); four mice were harvested per time point. We defined CT0 and 12 as the lights on (8am) and off (8pm) timings of the pre-DD light regime, respectively. Eight-week-old mice were used to ensure that the mice were within telogen phase when the epidermis was collected; any mouse in anagen phase was excluded.

### Generation of RE/RE mice

To generate RE/RE mice, *Bmal1*-stopFL mice (*6*) were crossed with K14-Cre mice (*46*) and Syt10-Cre mice (*20*). The resulting triple heterozygous mice (Arntl^stopFL/wt^; K14-Cre^tg/wt^;Syt10-Cre^tg/wt^) were then crossed with heterozygous *Bmal1*-stopFL mice (Arntl^stopFL/wt^; K14-Cre^wt/wt^;Syt10-Cre^wt/wt^) to generate the five RE genotypes used in this manuscript: i) WT (WT), Arntl^wt/wt^; K14-Cre^tg/wt^;Syt10-Cre^tg/wt^; ii) knockout (KO), Arntl^stopFL/stopFL^; K14-Cre^wt/wt^;Syt10-Cre^wt/wt^; iii) epidermis-RE (epi-RE), Arntl^stopFL/stopFL^; K14-Cre^tg/wt^;Syt10-Cre^wt/wt^; iv) brain-RE: Arntl^stopFL/stopFL^; K14-Cre^wt/wt^;Syt10-Cre^tg/wt^; and v) double reconstituted (RE/RE): Arntl^stopFL/stopFL^; K14-Cre^tg/wt^;Syt10-Cre^tg/wt^. All initial and maintenance crosses were performed using mice heterozygous for *Bmal1*-StopFL and each Cre, to avoid sterility associated with total BMAL1 loss or unwanted transgene integration effects, respectively. Only female mice expressing the Syt10-Cre were used for breeding due to the known expression of this Cre in the testes of male mice that may lead to general recombination of the floxed loci (*20*).

### Epidermis extraction

Mice were sacrificed by C0_2_ asphyxiation, and death was confirmed by cervical dislocation. The mice were then shaved, and the back skin was subsequently removed and placed in ice cold PBS. A tail sample was also taken to allow the genotype to be revalidated. To remove the hypodermis, the underside of the skin was then vigorously scraped using a scalpel. In a petri dish, the skin was then floated on 10 ml of 1 mg/ml dispase (Sigma Aldrich) in PBS for 45 min at 37°C, to separate enzymatically the epidermis. Subsequently, using a scalpel, the epidermis was scraped from the surface of the skin and then mechanically disaggregated using surgical scissors. The disaggregated epidermis was then placed into 10 ml of ice-cold PBS + 10% chelated FBS to inactivate the dispase. After vigorous shaking, the epidermal extract was sequentially filtered through 100 µM and 40 µM filters to give a purified extract of epidermal cells. Cell viability was checked via Trypan blue staining, and pellets of one million cells were then lysed in 500 µl of TRIzol (Invitrogen) for subsequent RNA extraction.

### Cell cycle analysis by flow cytometry

Epidermal cells were first extracted from 8-week-old male mice as described in ‘Epidermis extraction’ and then fixed in ice-cold 70% EtOH at -20°C for 2 hours. The cells were then washed 2x with PBS + 2% FBS + 1mM EDTA (wash buffer). The fixed cells were suspended to 1×10^7^cells/ml in wash buffer, and 100 µl of cell suspension was then mixed with 20 µl FITC anti-Ki67 antibody (BD Biosciences, 556026) and incubated at room temperature for 20 min. The cells were then washed 1x with wash buffer and suspended in 500 µl wash buffer + 5 µg/ml propidium iodide. The proportion of actively cycling cells (Ki67-positive) in G1, S and G2 phase of the cell cycle (cell DNA content: 1n – G1, > 1n and < 2n – S, 2n – G2) was interrogated using a using a Gallios flow cytometer (Beckman Coulter) and subsequent data analysis with FlowJo (v.10.8.0). Quantification was performed using a minimum of 10,000 cells/sample.

### RNA extraction

To extract total RNA, epidermal cells lysed in 500 µl TRIzol (Invitrogen) were vortexed with 100 µl chloroform, centrifuged for 15 min at 18,000 x*g* at 4°C, and then the upper aqueous layer containing RNA was isolated. The aqueous layer was then mixed with 825 µl RLT buffer (Qiagen) and 625 µl 100% ethanol to induce RNA precipitation. Samples were then processed using a RNeasy mini kit (Qiagen) according to the manufacturer’s protocol, and RNA was eluted in molecular biology–grade water and stored at –80°C.

### RNA sequencing

Total RNA from mice was quantified by Qubit® RNA BR Assay kit (Thermo Fisher Scientific) and the RNA integrity was estimated by using RNA 6000 Nano Bioanalyzer 2100 Assay (Agilent). RNA-seq libraries were prepared with KAPA Stranded mRNA-Seq Illumina® Platforms Kit (Roche) following the manufacturer’s recommendations. Briefly, following mRNA fragmentation, 500 ng of total RNA was used for the poly-A fraction enrichment with oligo-dT magnetic beads. Strand specificity was achieved for second-strand synthesis by using dUTP rather than dTTP. The blunt-ended, double-stranded cDNA was 3′-adenylated, and Illumina platform–compatible adaptors with unique dual indexes and unique molecular identifiers (Integrated DNA Technologies) were ligated onto them. The ligation product was enriched over 15 PCR cycles, and the final library was validated on an Agilent 2100 Bioanalyzer with the DNA 7500 assay. Libraries were sequenced using NovaSeq 6000 (Illumina) with a read length of 1×51bp+17bp+8bp, following the manufacturer’s protocol. Image analysis, base calling, and quality scoring of the run were processed using the manufacturer’s software Real Time Analysis (RTA 3.4.4).

Sequencing reads were first subjected to quality control and adapter trimming using Trimmomatic (version 0.36) (*47*). Reads were then aligned to the mm10 genome (UCSC) using the HISAT2 aligner (version 2.2.1) (*48*) and subsequently sorted using SAMtools (version 1.3.1) (*49*). After alignment, reads were quantified using featureCounts (version 1.6.0) (*50*), and the resulting count data was then normalized to generate RPKM values using the edgeR ‘rpkm’ function (version 3.18.1) (*51*).

### Identification of circadian genes and differential expression analysis

To generate consensus diurnal/circadian transcriptomes, the Jonckheere-Terpstra-Kendall (JTK_CYCLE) (*28*) and BIO_CYCLE (*27*) algorithms were used. In both instances, RPKM normalized gene expression values (log_2_ transformed) were used as input. Genes were considered to be rhythmically expressed if they showed a period of 24 hours and *P*-value ≤ 0.01. The lists of genes found to be rhythmically expressed were then intersected, and those genes identified as rhythmic by both algorithms were defined as the consensus diurnal/circadian transcriptome.

To identify specific genes reliant upon different clock communication pathways for their rhythmic expression, the DryR algorithm (*31*) was applied to all datasets. Raw RNAseq counts were used as input, and the function ‘dryseq()’ was applied to run the algorithm. The resulting output was then filtered (BICW ≥ 0.4, Amp ≥ 0.25, Periodicity = 24 hours), to generate the gene sets then further analyzed.

To identify genes that were differentially expressed irrespective of time-of-day, we applied the ‘deLimma’ function within the LimoRhyde (*52*) workflow to compare our WT and RE/RE diurnal RNA-seq datasets. Differentially expressed genes were defined as those having a Padj ≤ 0.05 and fold-change ≥ 1.5.

All heat maps were generated using the pheatmap package in the R programming environment. For heat maps of the consensus diurnal/circadian transcriptomes, the mean of RPKM normalized gene expression values (log_2_ transformed) from each time point were used as input to calculate a Z-score. For heat maps of the DryR-generated gene sets, the normalized read counts (as calculated by DryR) from each time point were used as input to calculate a Z-score. In all heat maps, genes were ordered by their calculated phase and a Z-score was calculated to visualize circadian expression patterns.

Amplitude and phase plots were generated using values calculated by the JTK_CYCLE algorithm (for plots related to the consensus diurnal/circadian transcriptomes) or DryR (for specific gene sets), and all graphs were made using the ggplot2 package in the R programming environment. To calculate amplitude density, the ‘geom_density’ function from the ggplot2 package was used to calculate a smoothed density estimate of the amplitude values.

### GO, Reactome, and motif analyses

All Gene Ontology (GO), Reactome and transcription factor target analyses were performed using the gene list enrichment analysis tool Enrichr (*53*). GO and Reactome terms found to be enriched within the lists ‘GO Biological Processes 2021’ and ‘Reactome 2022’, respectively, were analyzed further. Enriched terms shown in the figures of this paper were manually filtered to remove redundant terms and to emphasize the enrichment of biologically important categories. To identify enrichment for known transcription factor targets, the dataset ‘ENCODE and ChEA consensus TFs from ChIP-X’ was used. The complete output from Enrichr for GO and Reactome can be found in supplementary tables 1 and 2.

### Statistics

All information related to replicate inclusion/exclusion criteria, total number of replicates and other relevant statistical information is stated in the figure legends for all graphs.

**Supplemental table 1 – Specific gene sets generated by this study**

**Supplemental table 2 – Gene Ontology and Reactome analyses generated by this study**

## Supplemental information

### Supplementary text

A key finding of our manuscript is that reconstituting clock activity in the epidermis, brain or both tissues restores rhythmicity to certain elements of epidermal daily physiology. Mechanistically, clock communication may give rise to gene expression rhythmicity by: i) inducing rhythms in single cells; ii) inducing phase coherence between cells of a tissue; iii) inducing phase coherence between individuals. If inter-individual synchrony (iii) were being imposed, we would expect to observe a difference in gene expression standard deviation (due to a phase dispersion of biological replicates) between the different models, for commonly rhythmic genes. However, standard deviation of gene expression does not vary between RE/RE and WT epidermis for RE/RE+WT only rhythmic genes (Figure S3A, gene set shown in Figure 2A), or between brain-RE, RE/RE and WT for brain-RE+RE/RE+WT only rhythmic genes (Figure S3B, gene set shown in Figure S2E), indicating similar inter-individual synchrony in these conditions. Moreover, although generally (in all expressed genes, irrespective of rhythmicity) standard deviation of gene expression is significantly increased in RE conditions vs WT (Figure S3C), the change is minimal (3-6%), indicating that differences in biological variability do not meaningfully distinguish the models we analyse. In a previous manuscript, we also explored in depth these three mechanisms, but specifically in the context of the epi-RE model (1). In particular, we provided evidence that single-cell desynchrony (i) or intra-tissue desynchrony (ii) explained the reduced amplitude of epidermal core clock rhythmicity of epi-RE relative to WT mice. Together, we believe this makes imposing inter-individual synchrony (iii) an unlikely mechanism for the clock communication pathways we have analysed.

A mechanism imposing general intra-tissue synchrony (ii) would be predicted to increase the amplitude of rhythmic gene expression due to lessened phase dispersion of the individual cells. However, the amplitude of rhythmic gene expression is not significantly different between the consensus rhythmic transcriptomes of RE/RE and WT epidermis (Figure S3D), or their core clock genes (Figure S3E), suggesting comparable intra-tissue synchrony in both conditions. Furthermore, there is no residual rhythmicity in RE/RE of genes identified as rhythmic only in WT (Figure 2E). Together, this argues that imposing intra-tissue synchrony is not a mechanism that underlies peripheral:epidermal clock communication. Nonetheless, the consensus rhythmic transcriptome amplitude in epi-RE and brain-RE epidermis, and the amplitude of the epi-RE core clock, is significantly decreased relative to WT (Figure S3D and 1B). This indicates that brain:epidermis communication may be driving either single-cell synchrony (i) or intra-tissue synchrony (ii) to endow the restored physiological rhythmicity we observe in the RE/RE epidermis.

Imposing single-cell rhythmicity (i) can be achieved by a number of different mechanisms. One possibility is that clock communication ensures or optimises core clock function. However, the amplitude of all core clock, and key clock controlled genes, is indistinguishable between WT and RE/RE conditions (Figure S3E), indicating that enhancing core clock function via peripheral:epidermal clock communication is unlikely to explain why certain pathways are only rhythmic in WT epidermis. Moreover, although the epidermal core clock of epi-RE is dampened relative to WT and RE/RE (Figure 1B), genes identified by DryR as rhythmic in WT+RE/RE only (requiring brain:epidermal communication) show no apparent residual rhythmicity in epi-RE (Figure 2A). This suggests that whilst driving physiological activity of the epidermal clock may be a key element of brain:epidermal communication, additional mechanisms are also likely required.

To further explore potential mechanisms underlying clock communication, we used DryR to categorise each class of rhythmic genes by their expression magnitudes in each condition (e.g. rhythmic in X and Y, but with a lower mean expression in X etc.). The majority of genes requiring brain:epidermal communication (WT+RE/RE only, 629 genes - Figure 2A), segregated into two classes: i) Equivalent mean expression in all conditions (211 genes) (Figure S4A); ii) Equivalent mean expression in WT, RE/RE and epi-RE, but decreased or increased expression in brain-RE (205 genes) (Figure S4C). Strikingly, genes equivalently expressed in all conditions (i) were enriched for pathways related to the late cell cycle, indicating that rhythmicity of this process is restored by imposing rhythmicity upon genes already expressed in the cell (Figure S4B). Interestingly, genes in class (ii) with peak expression in the late night/early morning (ZT20-8) were generally upregulated at all time points in brain-RE, whilst those peaking in the afternoon/early night (ZT8-20) were generally repressed (Figure S4C). This pattern implies that a brain clock signal(s) is able to upregulate or repress the expression of these specific epidermal clock targets, dependent upon their expression phase. Although an interesting observation, we could not identify enrichment for specific pathways amongst this gene set (Figure S4D), and further work will be required to determine the nature and physiological significance of this signal(s).

Analysis of genes requiring peripheral:epidermal clock communication (WT only, 389 genes – Figure 2E) also identified two expression profiles describing the majority of genes: i) Equivalent mean expression in all conditions (166 genes) (Figure S4E); ii) Higher expression in WT relative to all other conditions (77 genes) (Figure S4G). Those with equivalent mean expression (i) were enriched for terms related to ribosome biogenesis (Figure S4F), but, in contrast, genes upregulated in WT (2) were enriched for extra cellular matrix terms (Figure S4H). This suggested that regaining rhythmicity of ribosome biogenesis required imposing rhythmicity upon already expressed genes, whilst for ECM-related processes a dual upregulation and gain of rhythmicity is required.

### Supplementary figure legends

**Figure S1.**
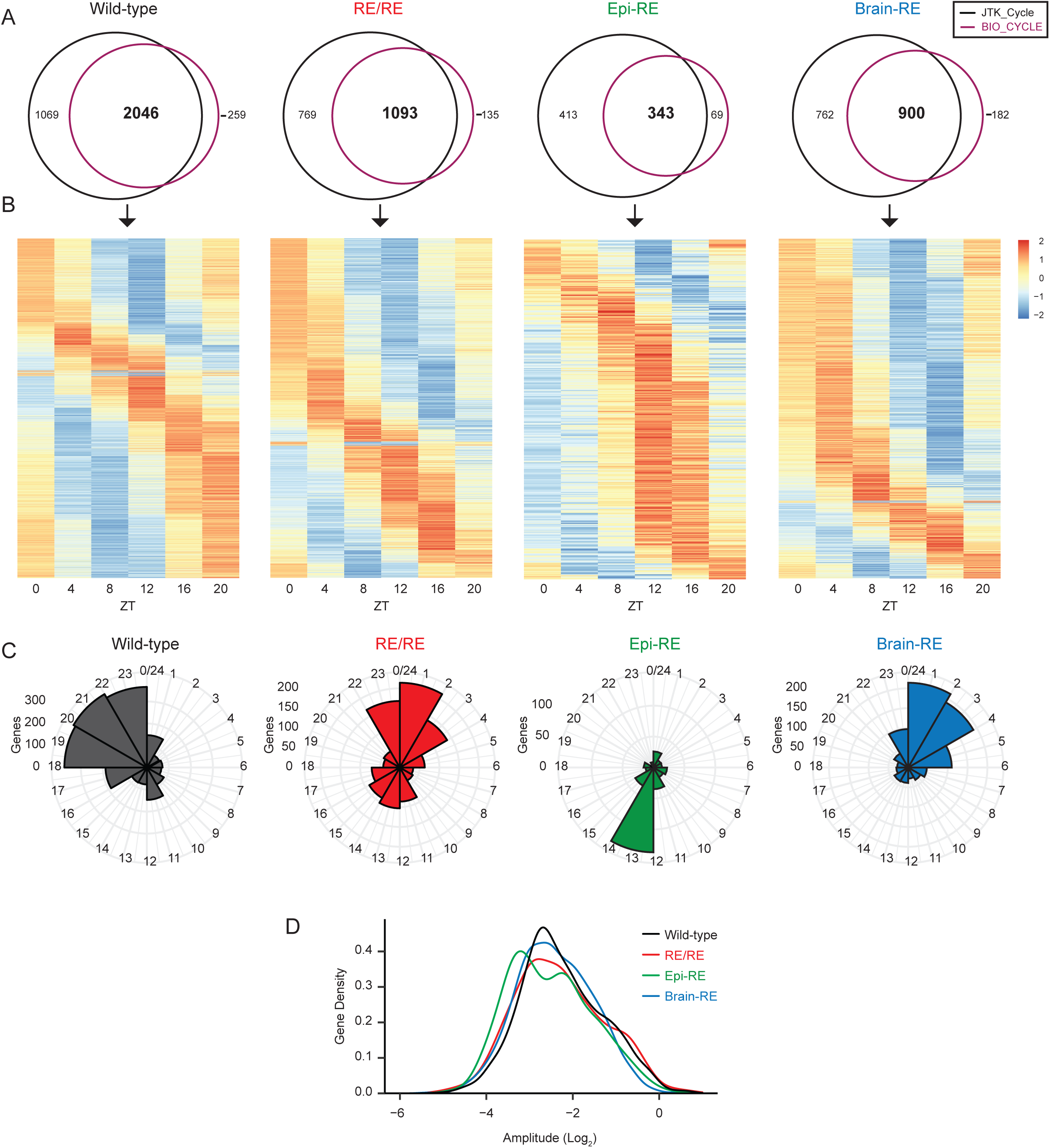
Tissue-specific reconstitution of *Bmal1* expression isolates communication between the epidermal clock and brain clock, related to Figure 1: (**A**) Venn diagrams quantifying the intersect of diurnal transcriptomes identified using the JTK_CYCLE and BIO_CYCLE algorithms for all clock reconstitution conditions. Rhythmicity was defined as a periodicity of 24 hours and a *P*-value ≤ 0.01 in both instances. (**B**) Heat map of the consensus diurnal transcriptomes (overlap of BIO_CYCLE and JTK_CYCLE-calculated diurnal transcriptomes) derived for each clock reconstitution condition (see A). Colors represent a Z-score calculated using the mean RPKM from each time point. Genes are sorted by expression phase. (**C** and **D**) Circular histograms (C) and a density plot (D) indicating the distribution of expression phases and expression amplitudes (respectively) of genes constituting the epidermal diurnal transcriptome of each clock reconstitution condition. Both expression phases and expression amplitudes are those calculated by JTK_Cycle. Brain-RE, Bmal1 reconstituted only in brain; Epi-RE, Bmal1 reconstituted only in epidermis; RE/RE, Bmal1 reconstituted in brain and epidermis.

**Figure S2.**
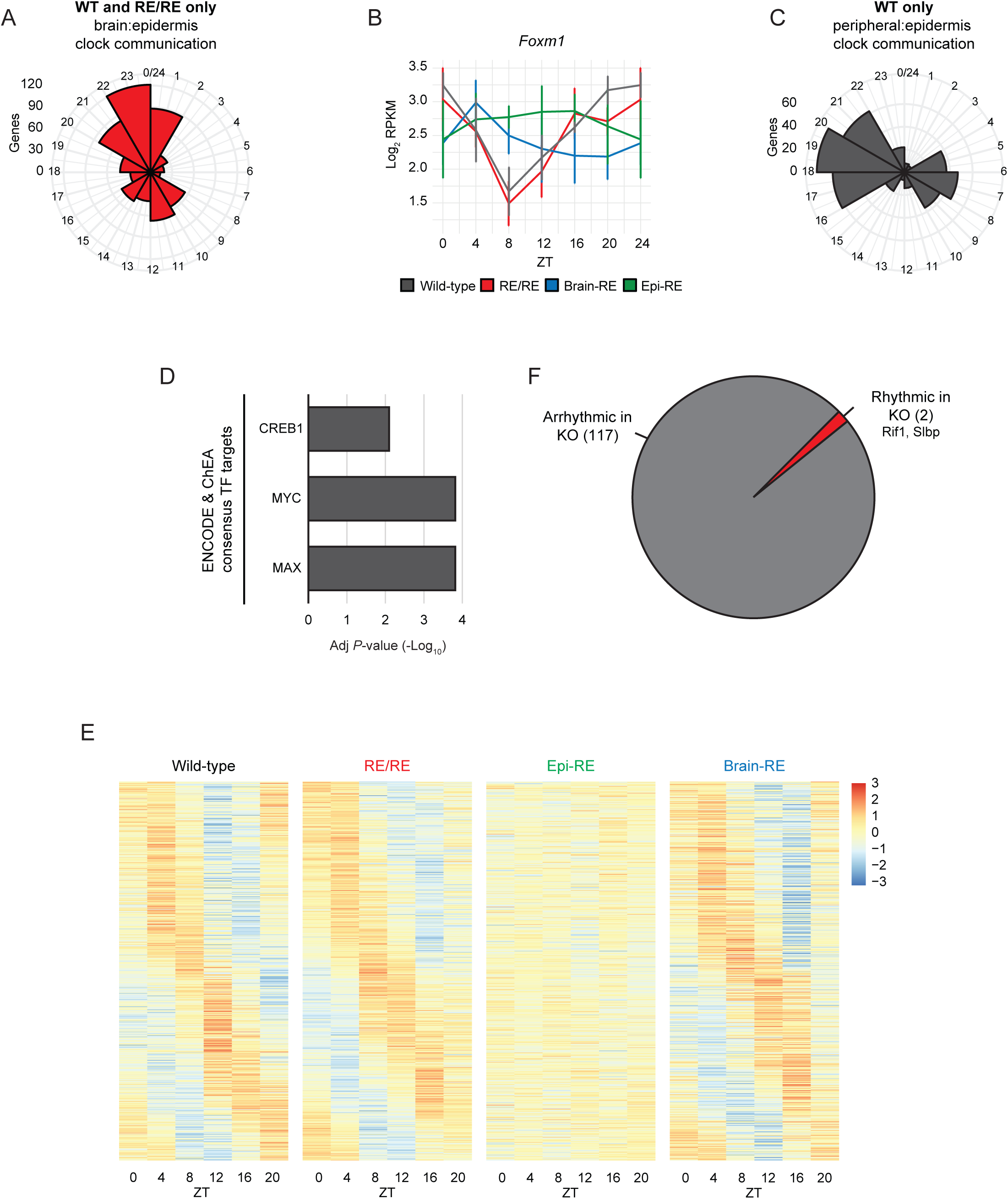
Communication with brain and other peripheral tissues ensures intact daily epidermal physiology, related to Figures 2 and 3: (**A** and **C**) Circular histograms indicating the distribution of expression phases of genes equivalently rhythmic only in WT and RE/RE epidermis (629 genes) (A) or only rhythmic in WT epidermis (399 genes) (C). The phase values plotted correspond to those calculated by DryR for this model, and therefore the phase of all genes in (B) is considered equivalent for WT and RE/RE epidermis. (**B**) Diurnal expression pattern of *Foxm1* in the epidermis of each clock reconstitution condition, as determined by RNA-seq. Values represent the log_2_-transformed mean (± standard deviation) RPKM of *n* = 4 biological replicates. The time point ZT0 is double-plotted for better visualization. (**D**) Enrichment of known transcription factor targets amongst genes rhythmic in WT epidermis only. **(E)** Heat maps showing the diurnal expression patterns of genes equivalently rhythmic in WT, RE/RE and brain-RE epidermis, but not epi-RE (608 genes), in all clock reconstitution conditions. Colors represent a Z-score calculated using the mean normalized counts (as calculated by DryR) from each time point. Genes are sorted by expression phase. **(F)** Pie chart indicating the number of the genes with antiphasic rhythmicity in brain-RE epidermis that also exhibited rhythmicity in the epidermis of Bmal1-StopFL mice (whole-body Bmal1 null). Data for Bmal1-StopFL mice were sourced from Welz *et al*, (2019), and rhythmicity was defined as a JTK_CYCLE calculated periodicity of 24 hours and *P* ≤ 0.01. Brain-RE, Bmal1 reconstituted only in brain; ChEA, ChIP Enrichment Analysis; ENCODE, Encyclopedia of DNA Elements; Epi-RE, Bmal1 reconstituted only in epidermis; RE/RE, Bmal1 reconstituted in brain and epidermis.

**Figure S3.**
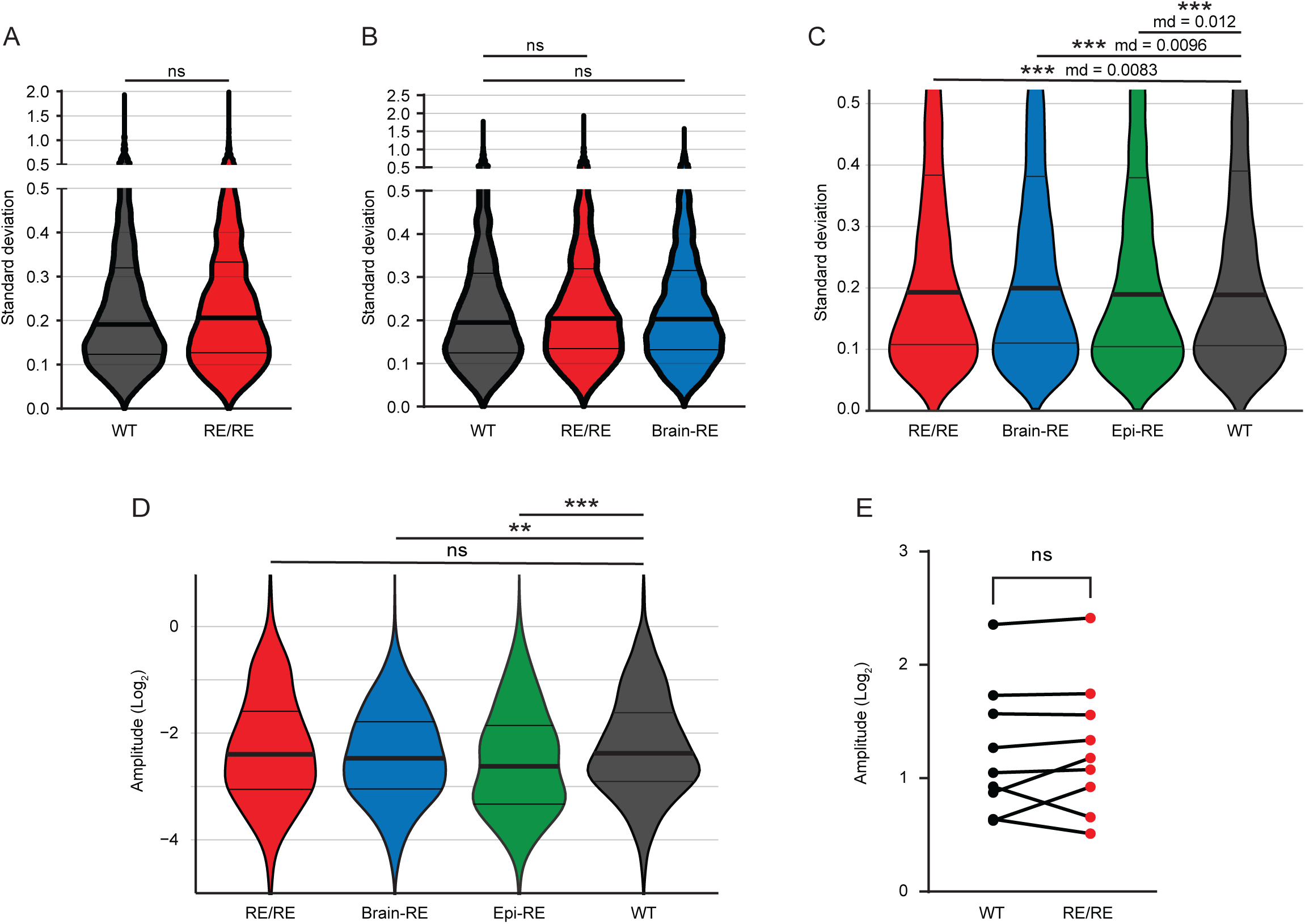
Gene expression arrhythmicity does not originate from inter-individual phase dispersion. **(A&B)** Violin plot showing the distribution of gene expression standard deviations for all time points of genes rhythmic in WT+ RE/RE epidermis only (629 genes - Figure 2A) (A) and WT+RE/RE+brain_RE epidermis only (608 genes - Figure S2E). Lines represent median +/- interquartile range. To calculate significance, a paired student t-test (A) or one-way ANOVA test (B) was performed. **C)** Violin plot showing the distribution of gene expression standard deviations for all time points of all genes in all conditions. Genes with a standard deviation of zero in any condition were excluded. Box plots represent median +/- interquartile range. Values > 1 are not displayed for visualisation purposes. To calculate significance, a one-way ANOVA test was performed. **D)** Violin plot showing the distribution of expression amplitudes for the consensus rhythmic transcriptomes of all conditions (Figure 1D). Box plots represent median +/- interquartile range. Values > 1 and < -5 are not displayed for visualisation purposes. To calculate significance, an unpaired student t-test was performed, **E)** A dot plot comparing the gene expression amplitude of core clock genes in WT and RE/RE epidermis. Each dot represents one of the core clock genes displayed in Figure 1B. To calculate significance, a paired student t-test was performed. Md, mean difference vs WT; ns, non-significant; **P < 0.01; ***P < 0.001.

**Figure S4.**
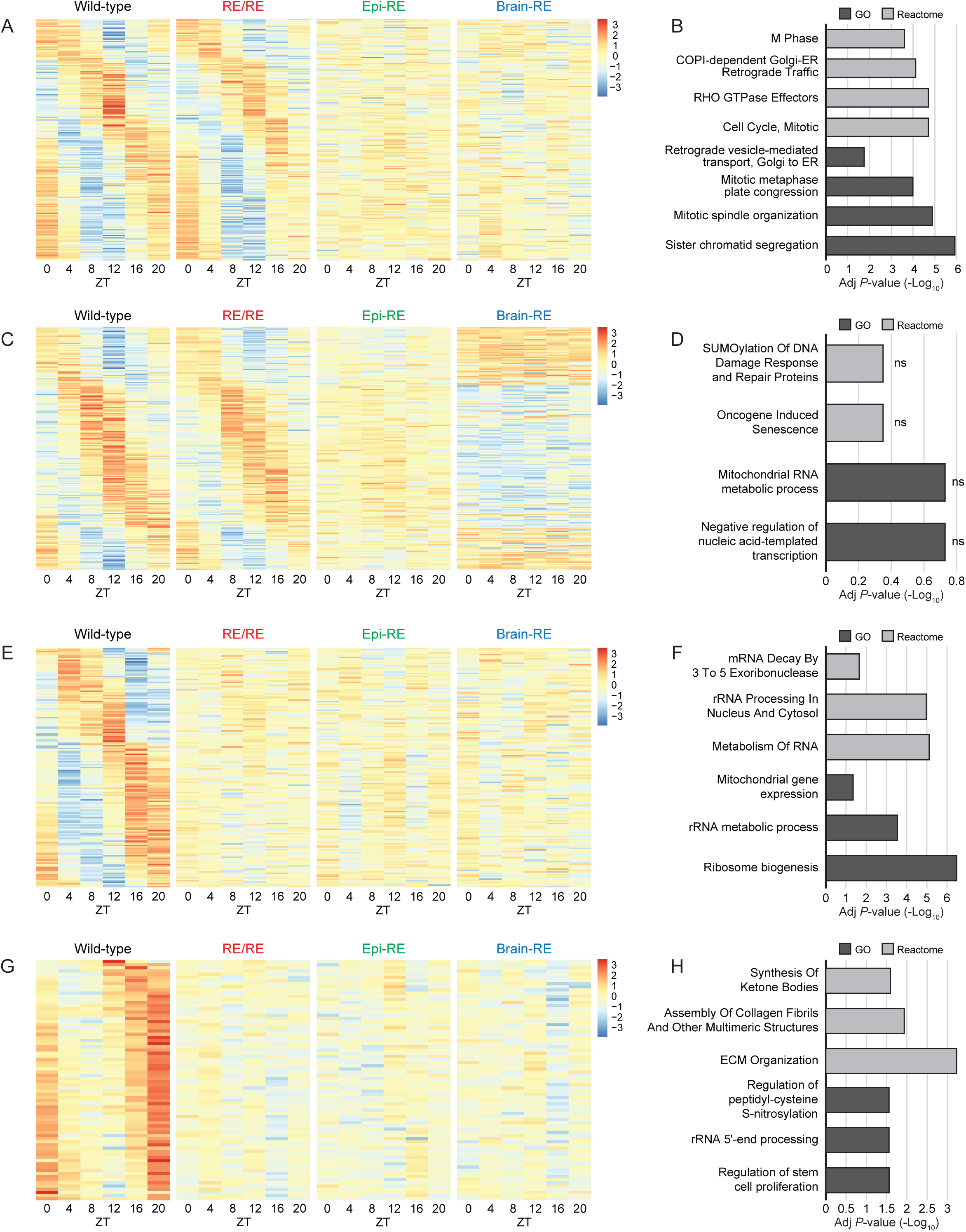
Expression magnitude classification of genes regulated by different clock communication pathways. **(A, C, E and G)** Heat maps showing the diurnal expression patterns of: (A) genes requiring brain:epidermal communication (WT+RE/RE only, 608 genes - Figure 2A) that also show equivalent mean expression in all conditions (211 genes); (C) Genes requiring brain:epidermal communication (WT+RE/RE only, 608 genes - Figure 2A) that show equivalent mean expression in WT, RE/RE and epi-RE, but decreased or increased expression in brain-RE (205 genes); (E) Genes requiring peripheral:epidermal clock communication (WT only, 389 genes – Figure 2E) that show equivalent mean expression in all conditions (166 genes); (G) Genes requiring peripheral:epidermal clock communication (WT only, 389 genes – Figure 2E) that show higher expression in WT relative to all other conditions (77 genes). Genes were categorized by their most probable ‘mean model’ as determined by DryR. Colors represent a Z-score calculated using the mean normalized counts (as calculated by DryR) from each time point. Genes are sorted by expression phase. **(B, D, F and H)** Selected Gene Ontology and Reactome terms enriched amongst the gene classes presented in the heatmap shown opposite.

**Figure S5.**
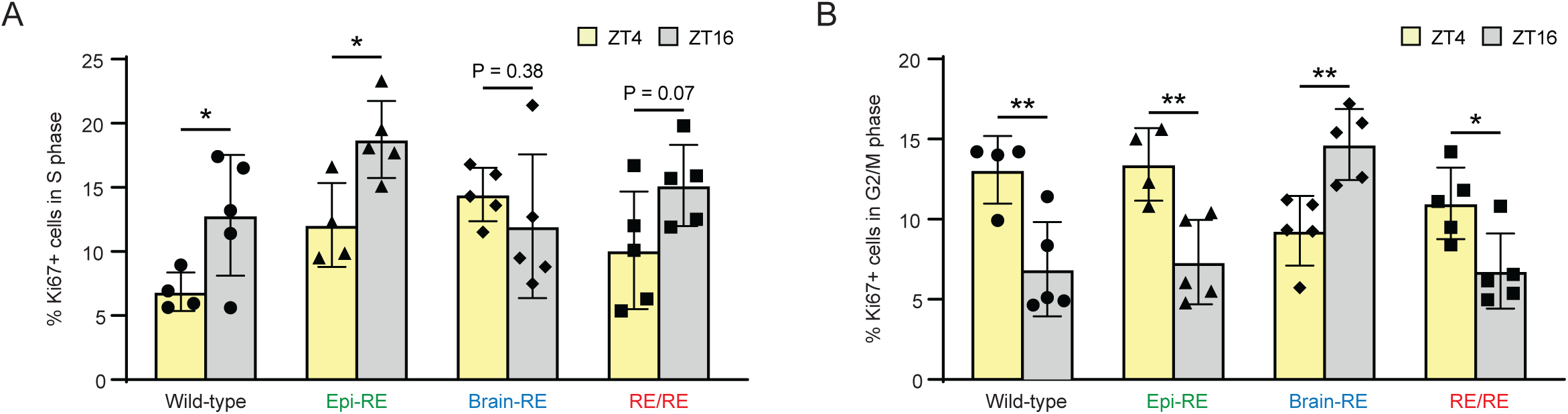
Brain clock signals drive an antiphasic pattern of epidermal DNA replication, related to Figure 4: (**A** and **B**) Percentage of actively cycling epidermal cells (Ki67-positive) in S (> 1n - < 2n DNA content) (B) or G2/M (2n DNA content) phase (C) at ZT4 and ZT16, in 35-week-old female mice. Values represent the mean (± standard deviation) of *n* = 4-5 biological replicates. \**P* ≤ 0.05, \*\**P* ≤ 0.01. Brain-RE, Bmal1 reconstituted only in brain; Epi-RE, Bmal1 reconstituted only in epidermis; RE, reconstituted; RE/RE, Bmal1 reconstituted in brain and epidermis; ZT, Zeitgeber time.

**Figure S6.**
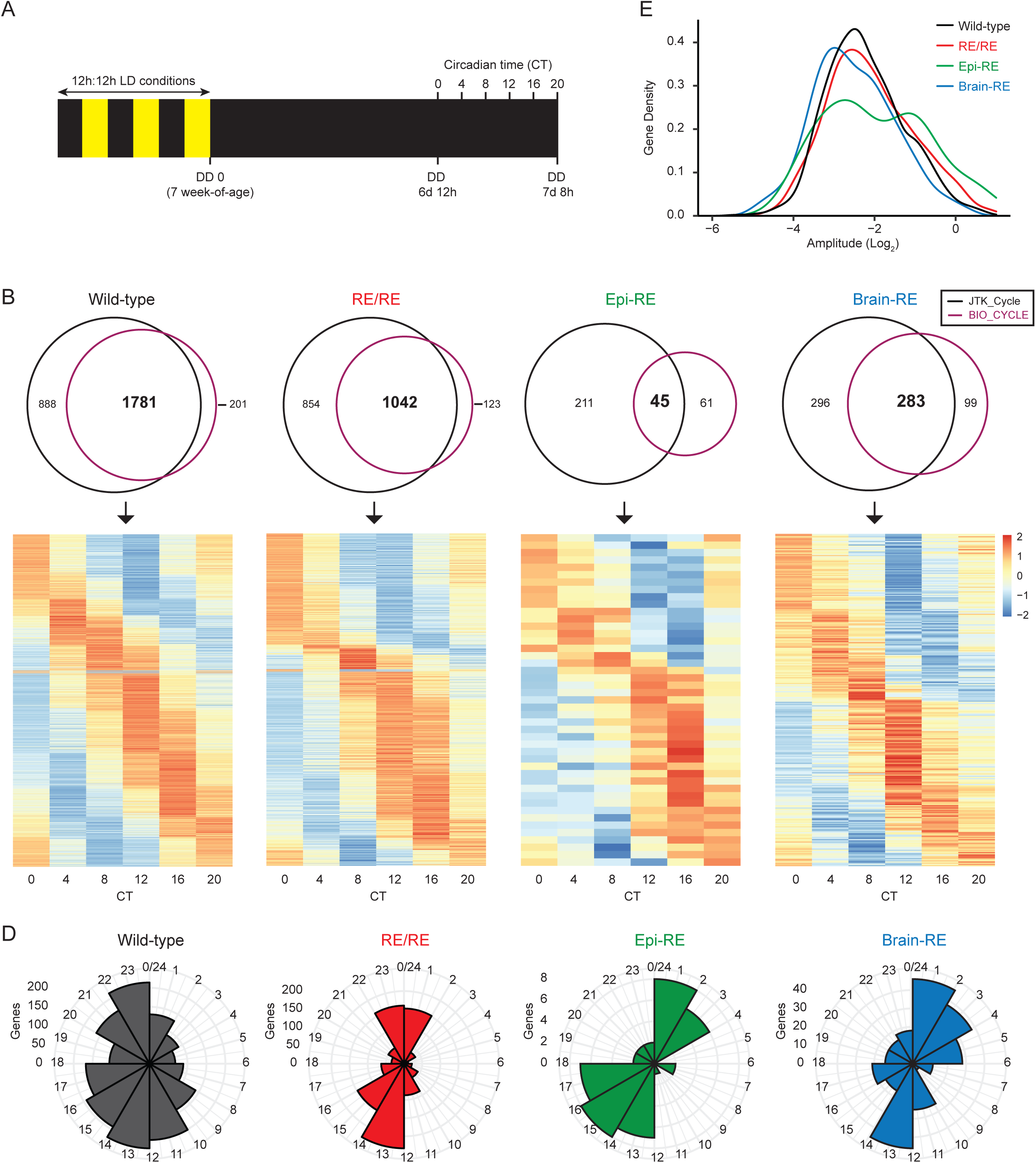
Brain clock communication ensures robust daily epidermal physiology in constant darkness, related to Figure 5: (**A**) Venn diagrams quantifying the intersect of circadian transcriptomes identified by the JTK_CYCLE and BIO_CYCLE algorithms for all clock reconstitution conditions. Rhythmicity was defined as a periodicity of 24 hours and a *P*-value ≤ 0.01 in both instances. (**B**) Heat map of the consensus circadian transcriptomes (overlap of BIO_CYCLE and JTK_CYCLE-calculated circadian transcriptomes) derived for each clock reconstitution condition (see A). Colors represent a Z-score calculated using the mean RPKM from each time point. Genes are sorted by expression phase. (**C** and **D**) Circular histograms (C) and a density plot (D) indicating the distribution of expression phases and expression amplitudes (respectively) of genes constituting the epidermal circadian transcriptome of each clock reconstitution condition. Both expression phases and expression amplitudes are those calculated by JTK_Cycle. Brain-RE, Bmal1 reconstituted only in brain; Epi-RE, Bmal1 reconstituted only in epidermis; RE/RE, Bmal1 reconstituted in brain and epidermis.

**Figure S7.**
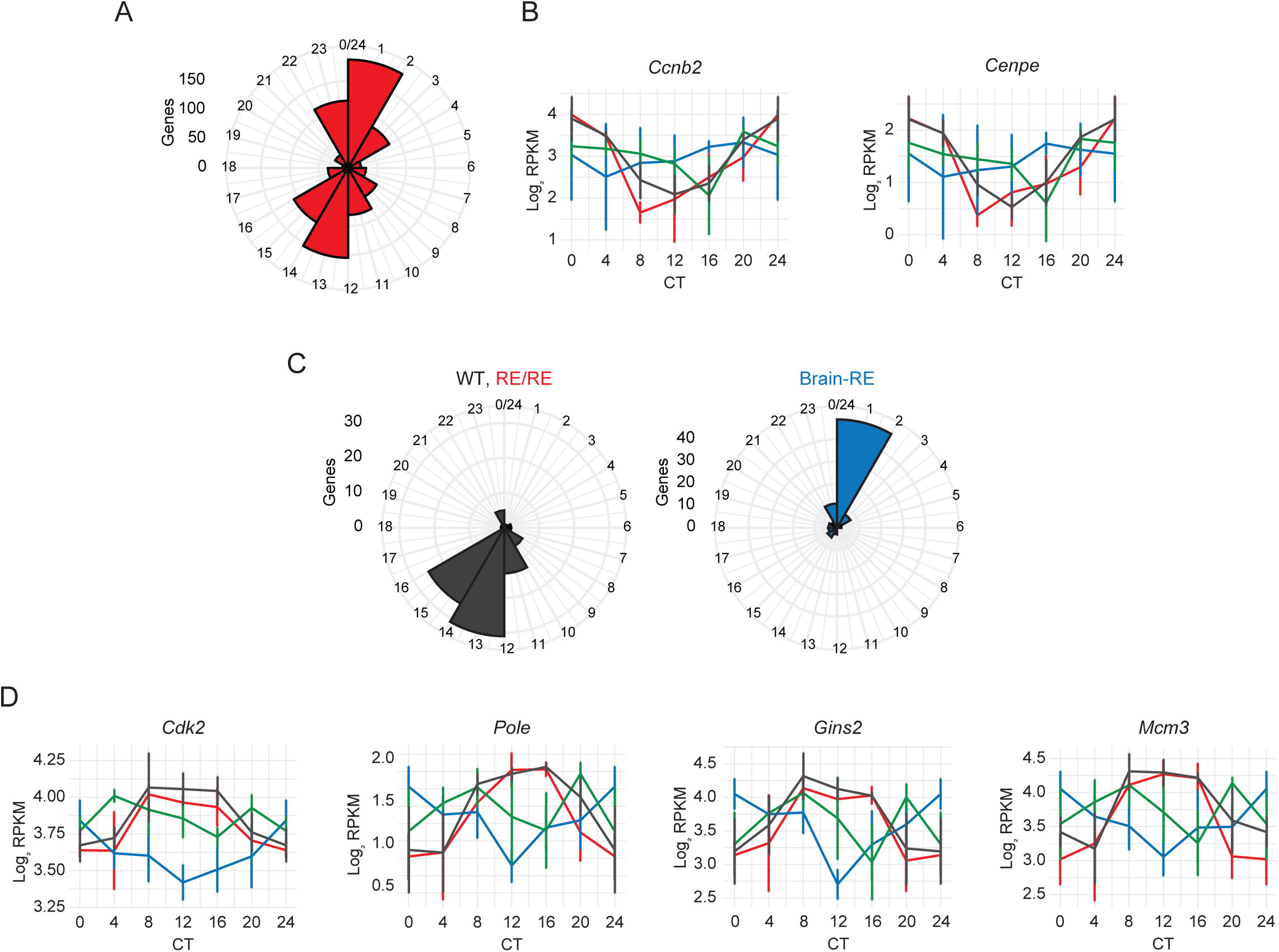
Brain clock communication ensures robust daily epidermal physiology in constant darkness, related to Figure 5: (**A** and **C**) Circular histograms indicating the distribution of expression phases of genes equivalently rhythmic only in WT and RE/RE epidermis (961 genes) (A) or with antiphasic rhythmicity in brain-RE epidermis relative to wild-type and Brain-RE (89 genes) (C). The phase values plotted in (C) correspond to those calculated by DryR for this model, and therefore the phase of all gene is considered equivalent in wild-type and Brain-RE. (**B** and **D**) Circadian expression patterns of late (B) and early (D) cell cycle regulators in the epidermis of each clock reconstitution condition, as determined by RNA-seq. Values represent the log_2_-transformed mean (± standard deviation) RPKM of *n* = 4 biological replicates. The time point CT0 is double-plotted for better visualization. Brain-RE, Bmal1 reconstituted only in brain; Epi-RE, Bmal1 reconstituted only in epidermis; RE/RE, Bmal1 reconstituted in brain and epidermis.

